# The novel *Dbl* homology/BAR domain protein, MsgA, of *Talaromyces marneffei* regulates yeast morphogenesis during growth inside host cells

**DOI:** 10.1101/481408

**Authors:** Harshini Weerasinghe, Hayley E. Bugeja, Alex Andrianopoulos

## Abstract

Microbial pathogens have evolved many strategies to evade recognition by the host immune system, including the use of phagocytic cells as a niche within which to proliferate. Dimorphic pathogenic fungi employ an induced morphogenetic transition, switching from multicellular hyphae to unicellular yeast that are more compatible with intracellular growth. A switch to mammalian host body temperature (37°C) is a key trigger for the dimorphic switch. This study describes a novel gene, *msgA*, from the dimorphic fungal pathogen *Talaromyces marneffei* that controls cell morphology in response to host cues rather than temperature. The *msgA* gene is upregulated during murine macrophage infection, and deletion results in aberrant yeast morphology solely during growth inside macrophages. MsgA contains a Dbl homology domain, and a <u>B</u>in, <u>A</u>mphiphysin, <u>R</u>vs (BAR) domain instead of a Plekstrin homology domain typically associated with guanine nucleotide exchange factors (GEFs). The BAR domain is crucial in maintaining yeast morphology and cellular localisation during infection. The data suggests that MsgA does not act as a canonical GEF during macrophage infection and identifies a temperature independent pathway in *T*. *marneffei* that controls intracellular yeast morphogenesis.

## Introduction

Host defence against disease causing microbes involves actively identifying and eliminating invading pathogens. This begins with the innate immune system where host phagocytes engulf and destroy these microbes, followed by triggering of an adaptive immune response. The success of a pathogen depends on its ability to escape the host response and for some pathogens this involves residing within particular phagocytic cells of the host where they are then able grow and proliferate. An important factor facilitating this lifestyle is the capacity to adopt a growth form suitable for the spatial constraints of the intracellular environment of a host cell. For many fungal pathogens that utilise the intracellular environment of macrophages as a means of immune system avoidance, morphological plasticity is an important virulence attribute (Nemecek et al. 2006; Nguyen & Sil 2008; Webster & Sil 2008; Beyhan et al. 2013; Brandhorst et al. 1999; Rooney et al. 2001). This is exemplified by the dimorphic fungi, which display saprophytic, multicellular, filamentous hyphal growth in the external environment and are able to adopt a unicellular yeast growth form during infection. Dysregulation of morphology, leading to the production of a growth form that disrupts the integrity of the phagocytic cells, exposes the pathogen to the entire immune system.

*Talaromyces marneffei* (formerly *Penicillium marneffei*) is an intracellular human pathogen that exhibits temperature dependent dimorphic growth. At 25°Cit grows in a multinucleate septate hyphal form that can differentiate to produce uninucleate asexual spores (conidia) (Andrianopoulos 2002). At 37°C*in vitro* it grows as uninucleate yeast that divides by fission. The transition from hyphal cell or dormant conidium to a yeast cells proceeds by arthroconidiation, a process of filament fragmentation, in which nuclear and cellular division are coupled, double septa are laid down with subsequent cell separation producing yeast cells (Andrianopoulos 2002). The transition from yeast cell to hyphal cell occurs by polarisated growth at the tips of the elongate yeast cells and a switch to cell division by septation without subsequent cell separation.

The conidia are the infectious propagules and infection is believed to be initiated through their inhalation (Vanittanakom et al. 2006). Once within the human host, *T*. *marneffei* conidia are engulfed by host primary alveolar macrophages where they bypass the process of arthroconidiation and germinate directly into yeast cells, which are the pathogenic form. While hyphae and yeast are the predominant morphologies for *T*. *marneffei*, particular growth conditions manifest distinct differences in cell shape and length for some cell types. For instance, *in vitro* grown *T*. *marneffei* yeast cells have an elongated, filament-like morphology, whilst *in vivo* produced yeast cells are short, ellipsoid and compact, a form more suited for growing within the confines of the macrophage. As an intracellular pathogen the ability of *T*. *marneffei* to tightly regulate the dimorphic switch and maintain yeast morphology in the host is crucial for pathogenicity. Therefore, temperature drives the dimorphic switch and host signals modify yeast cell morphogenesis.

In a number of animal and plant pathogenic fungal species, cell type specific morphogenesis is controlled by signalling pathways involving the small GTPase molecular switches and their accessory factors; guanine-nucleotide exchange factors (GEFs) and GTPase-activating proteins (GAPs) (reviewed in (Chant & Stowers 1995)). These play crucial roles in processes associated with cell shape such as control of polarized growth and cytoskeletal organisation. Rho-type GEFs are responsible for activating their small GTPase targets and their activity has been shown to regulate cytokinesis, cell wall integrity, antifungal resistance, viability, tissue invasive growth, mycotoxin production and virulence (Fuchs et al. 2007; Tang et al. 2005; Herrmann et al. 2014; Zhang et al. 2018). The majority of these studies have focused on GEFs containing canonical structural domains - a catalytic *<u>D</u>bl* <u>h</u>omology domain (DH) domain and an auxiliary plekstrin homology (PH) domain, excluding the contribution of GEFs with alternate domain structures (reviewed in Rossman et al. 2005).

In *T*. *marneffei*, a number of small GTPases of the Ras superfamily and associated downstream effectors have been shown to influence morphogenesis of all cell types in specific and distinct patterns, during growth under both *in vitro* and *ex vivo* (intracellular macrophage) growth conditions, by affecting polarity establishment and the differentiation of distinct cell types (Boyce & Andrianopoulos 2011). For example, the Cdc42 orthologue in *T*. *marneffei*, encoded by *cflA*, is required for correct yeast and hyphal cell morphogenesis *in vitro*, as well as germination of conidia but does not affect the various cell types or differentiation of asexual development structures (conidiophores). Whereas the Rac orthologue, encoded by *cflB*, is important for hyphal cell morphogenesis and asexual development but not yeast cell morphogenesis (Boyce et al. 2001; Boyce et al. 2003). In addition, orthologues of the p21 activated kinases (PAK) Ste20 and Cla4, encoded by *pakA* and *pakB* respectively, are important for conidial germination and yeast cell morphogenesis at 37°C(Boyce & Andrianopoulos 2007; Boyce et al. 2009). In particular *pakB* exclusively affects the formation of yeast cells during intracellular growth, suggesting that the pathways controlling morphogenesis of *T*. *marneffei in vitro* and during intracellular growth in host cells have some unique components that respond to distinct cues.

In a *T*. *marneffei* expression profiling study examining genes expressed during growth inside macrophages, a gene was identified that was specifically upregulated inside murine macrophages and was predicted to encode a RhoGEF-like protein (Weerasinghe et al 2018 *in prep*). This previously uncharacterised gene that was named *msgA* (<u>m</u>acrophage <u>s</u>pecific G<u>EF-like</u> A) is the only RhoGEF-like encoding gene in *T*. *marneffei* to show this specific pattern of expression. While MsgA contains a DH domain, unlike canonical GEFs, a Bin-Amphiphysin-Rvs (BAR) domain replaces the PH domain. Proteins containing BAR domain have been shown to play various roles in membrane dynamics as well as interacting with small GTPases and many other proteins (Habermann 2004; Zimmerberg & McLaughlin 2004; Dawson et al. 2006; Peter et al. 2004). Deletion of *msgA* results in aberrant yeast morphology during macrophage infection but not during *in vitro* growth at 37°C. Induced overexpression of *msgA* during *in vitro* growth resulted in yeast cell formation mimicking that of growth inside macrophages. Mutational analysis showed that the BAR domain of MsgA is crucial in establishing correct yeast morphogenesis and localisation during intracellular growth. Together these results define a novel host infection specific pathway that regulates intracellular morphogenesis in *T*. *marneffei*.

## Materials and methods

### Molecular methods and plasmid construction

*T*. *marneffei* genomic DNA was isolated as previously described (Borneman et al. 2000). Southern and northern blotting was performed using Amersham Hybond N+ membrane and [a^-32^P]dATP labelled probes by standard methods (Sambrook et al. 1989). Sequences of primers are provided in Supplementary Table 1.

Deletion constructs were created using a modified Gateway™ method (Boyce et al. 2012). The deletion construct of *msgA* was created using pHW7711 containing the pDONR-*pyrG* cassette. Wildtype PCR product of *msgA* (PMAA_089500) was generated with primers VV55 and VV56 and cloned into pBluescript II SK+ to generate pHW7897. To make the deletion construct, pHW7905, PCR primers VV57 and VV58 were used to generate an inverse PCR product for the Gateway™ reaction. To create a complementation construct, wildtype PCR product of *msgA* was cloned into the *niaD* targetting plasmid pHB7615 to generate pHW8053.

The overexpression allele *xylP(p)*::*msgA* (pHW8056) was generated by PCR amplification of 2 kb of the 5’ promoter and ORF regions of the *msgA* gene using WW78 and WW77 to create subclone pHW8069. An inverse PCR product of pHW8069 was generated by amplification with primers WW79 and WW42 that excluded the promoter region. Subsequently the *xylP(p)* fragment PCR amplified from pHW8056 using H57 and H56 was phosphorylated and ligated to the inverse PCR fragment from pHW8069.

To generate the *msgA*::*mCherry* construct, pHW8053 was inverse PCR amplified using WW46 and WW57 and the resultant product was cut with *XbaI*. The *mCherry* fragment from pHW7911 was isolated by digestion with *SpeI/EcoRV* and this was ligated into the pHW8053 inverse PCR product to generate pHW7965.

To generate the *msgA*^*ΔDH*^ and *msgA*^*ΔBAR*^ domain deletion alleles, pHW8053 was inverse PCR amplified using WW59 and WW60 (ΔDH), and WW44 and WW45 (ΔBAR), and the resulting products were phosphorylated and self-ligated to produce pHW7966 and pHW7964 respectively. To generate the *msgA*^*ΔDH*^*::mCherry* and *msgA*^*ΔBAR*^*::mCherry* the same primers were used to inverse PCR from pHW7965 to generate pHW7962 and pHW7963.

### Fungal strains and media

DNA-mediated transformation of *T*. *marneffei* was performed as previously described (Borneman et al. 2000). Strains used in this study are listed in Supplementary Table 2. Strains G816 (*ΔligD::pyrG*^-^*, niaD1*) and G809 (*ΔligD::pyrG*^*+*^*, niaD1*) were used as recipient strains to generate *ΔmsgA* and *xylP(p)::msgA* strains respectively (Bugeja et al. 2012). The *ΔmsgA* strain was generated by transforming G816 with the *XhoI/NotI* fragment of pHW7905 to delete *msgA* from −200 to +6040 (relative to the translational start). Transformants were selected for uracil prototrophy. The *xylP(p)::msgA* strain was generated by transforming G809 with the *NruI*/*EcoRV* fragment of pHW8056. Transformants were selected for glufosinate resistance. Strains bearing the *msgA*^*+*^, *msgA::mCherry*, *msgA*^ΔBAR^, *msgA*^ΔDH^, *msgA*^ΔBAR^*::mCherry* and *msgA*^ΔDH^*::mCherry* alleles in a Δ*msgA* background were generated by transforming G1003 with pHW8055, pHW7965, pHW7966, pHW7964, pHW7962 and pHW7963 respectively. These constructs were targeted to the *niaD* locus to generate strains G1045, G1046, G1047, G1048, G1049 and G1050 respectively. Homologous integration at *niaD* repairs the mutation in this strain, and transformants were selecting for their ability to utilise nitrate as a sole nitrogen source.

For induction of the *xylP* promoter during hyphal growth at 25°C*in vitro*, strains were grown in BHI medium with (inducing) or without (non inducing) 1% xylose for 4 days before microscopic observation. For growth in continuous inductive or non-inductive conditions, growth in each medium was extended for a further two days (total 6 days). For yeast growth at 37°C*in vitro*, strains were grown in BHI medium for 4 days then transferred to BHI medium with (inducing) or without (non inducing) 1% xylose for a further 2 days. For growth in continuous inductive or non-inductive conditions, strains were grown for 6 days in BHI with or without 1% xylose respectively.

In order to determine the septal span in hyphal cells for the wildtype and *xylP(p)*::*msgA* during induced *msgA* expression, the distance between septa was measured for a 100 subapical cellular compartments for both strains. Additionally to determine the branching frequency for the wildtype and *xylP(p)*::*msgA* during induced *msgA* expression, the number of branch points along a 100μm section of subapical hyphal cells for ten hyphae was counted for both strains.

### Preparation of RNA

RNA was isolated from two separate conditions for RT-PCR analysis. For the *in vitro* expression experiments RNA was isolated from FRR2161 (wildtype) yeast cells grown at 37°Cfor 6 days in liquid Brain Heart Infusion medium (BHI). For the macrophage infection expression experiments RNA was isolated from FRR2161 (wildtype) and *ΔmsgA* (G1003) cells isolated from infected lipopolysaccharide (LPS)(Sigma) activated J774 murine macrophages. J774 murine macrophages were seeded at a concentration of 1 x 10^6^/ml into 175cm^3^ sterile cell culture flasks in 20 mL of complete Dulbecco’s Modified Eagle Medium (DMEM, 10% foetal bovine serum, 8 mM L-glutamine and 1% penicillin-streptomycin), incubated at 37°Cfor 24 h and then activated with 0.1 μg/mL LPS for 24 h. Macrophages were washed in phosphate buffered saline (PBS) and 20 mL of complete DMEM containing 1 x 10^7^ conidia was added. Macrophages were incubated for 2 h at 37°C(to allow conidia to be phagocytosed), washed once in PBS (to remove free conidia) and incubated a further 24 h at 37°C. Macrophages were then treated with 4 ml of 0.25% v/v Triton X solution for 2 minutes at room temperature to lyse the macrophages and extract *T*. *marneffei*. Both the lysate and *T*. *marneffei* grown in DMEM medium without macrophages were centrifuged at 2000 rpm for 5 minutes at 4°C. The resultant pellets were washed in PBS and the RNA was extracted using TRIzol Reagent (Invitrogen) and a MP FastPrep-24 bead beater according to the manufacturer’s instructions. RNA was DNase treated (Promega) prior to expression analysis.

### Macrophage infection assay

J774 murine or THP-1 human macrophages were seeded at a concentration of 1 x 10^5^/ml into a 6 well microtitre tray containing one sterile coverslip and 2 mL of complete DMEM (J774) or RPMI (THP-1) per well. J774 macrophages were incubated at 37°Cfor 24 h before activation with 0.1 μg/mL LPS. THP-1 macrophages were differentiated with 32μM phorbol 12-myristate 13-acetate (PMA) at 37°Cfor 24 h. Macrophages were incubated a further 24 h at 37°C, washed in PBS and 2 mL of DMEM or RPMI medium containing 1 x 10^6^ conidia was added. A control lacking conidia was also performed. Macrophages were incubated for 2 h at 37°C(to allow conidia to be engulfed), washed once in PBS (to remove free conidia) and incubated a further 24 or 48 h at 37°Cafter the addition of 2 mL of fresh DMEM or RPMI medium. Macrophages were fixed in 4% paraformaldehyde and stained with 1 mg/mL fluorescent brightener 28 (calcofluor, CAL) to observe fungal cell walls. Mounted coverslips were examined using differential interference contrast (DIC) and epifluorescence optics for cell wall staining and viewed on a Reichart Jung Polyvar II microscope. Images were captured using a SPOT CCD camera (Diagnostic Instruments Inc) and processed in Adobe Photoshop™. The number of ungerminated conidia, germlings, filaments or yeast cells was recorded in a population of approximately 100 macrophages in three independent experiments. Filaments were defined as any fungal cell with at least one septum. Ellipsoid and elongated fungal cells with no septa were counted as yeast cells. Mean and standard error of the mean values were calculated.

### Microscopic techniques for morphological analysis of T. marneffei

For morphological and general growth examination, *T*. *marneffei* was grown in liquid BHI (Oxoid) medium at 25°Cfor 2 days (hyphal growth) or 37°Cfor 4 days followed by transfer of approximately 10% of the culture to fresh medium for a further 2 days at 37°C(yeast growth). Aliquots of 500μL were transferred to 1.5 mL microfuge tubes containing 20 μL of 4% paraformaldehyde in PME (50 mM piperazine-N, N’-bis(2-ethanesulfonic acid (PIPES), 1 mM MgSO4, 20 mM EGTA pH 6.7) fixing solution. Samples were incubated for 20 min at room temperature and then pelleted by centrifugation for 5 min. The supernatant was removed and pellets were resuspended in 0.001% v/v Tween 80 supplemented with fluorescent brightener 28 (final concentration 0.014 mg/mL) and observed microscopically.

### Immunofluorescence microscopy

Immunofluoresence localization of the *msgA::mCherry*, *msgA*^ΔDH^*::mCherry* and *msgA*^ΔBAR^*::mCherry* strains was performed with anti-mCherry rat monoclonal primary antibody (Life Technologies) and an anti-rat ALEXA 488 conjugated goat secondary antibody (Molecular Probes) using standard protocols (Fischer & Timberlake 1995). Immunofluoresence microscopy controls using only primary or secondary antibodies as well as an untagged strain were performed to confirm specificity of the antibodies.

### Germination tests

For *in vitro* germination experiments, approximately 10^5^ spores were inoculated into 200 µl of Synthetic Dextrose (SD) medium containing 10 mM (NH4)2SO4 and incubated for 8, 16, or 24 h at 37°C. The rates of germination were measured microscopically by counting the numbers of germinating conidia (conidia with a visible germ tube) in a population of approximately 100 fungal cells. Three independent experiments were performed. Mean and standard error of the mean values were calculated.

## Results

### The msgA gene encodes a unique RhoGEF-like protein with a distinctive domain structure and expression

A previous RNAseq study identified the *msgA* gene as showing specific transcriptional upregulation during intracellular growth of *T*. *marneffei* in J774 murine macrophages, relative to *in vitro* yeast or hyphal growth (Weerasinghe *et al*., in prep). Based on this expression profile and its domain structure the gene was denoted *msgA* (<u>m</u>acrophage <u>s</u>pecific <u>GEF-like</u>). The *msgA* gene is one of six genes in the *T*. *marneffei* genome predicted to encode RhoGEF-like proteins. These are CtlA (Cdc24 orthologue), TusA (Tus1 orthologue), RomB (Rom1/Rom2 orthologue), BudC (Bud3 orthologue) and RgfF (<u>R</u>ho<u>GEF</u> 6), based on the annotations in *Saccharomyces cerevisiae* (Cdc24, Tus1 and Rom1/Rom2), *Aspergillus nidulans* (Bud3) or having been previously undesignated (*msgA* and *rgfF*) (Figure 1A). RhoGEF proteins from a number of dimorphic and/or pathogenic fungi were examined and while orthologues of MsgA exist in many phyla they are absent from the Saccharomycotina sub-phylum and Basidiomycete phylum. It is also clear that MsgA is part of a distinct clade of RhoGEF-like proteins.

**Figure 1.**
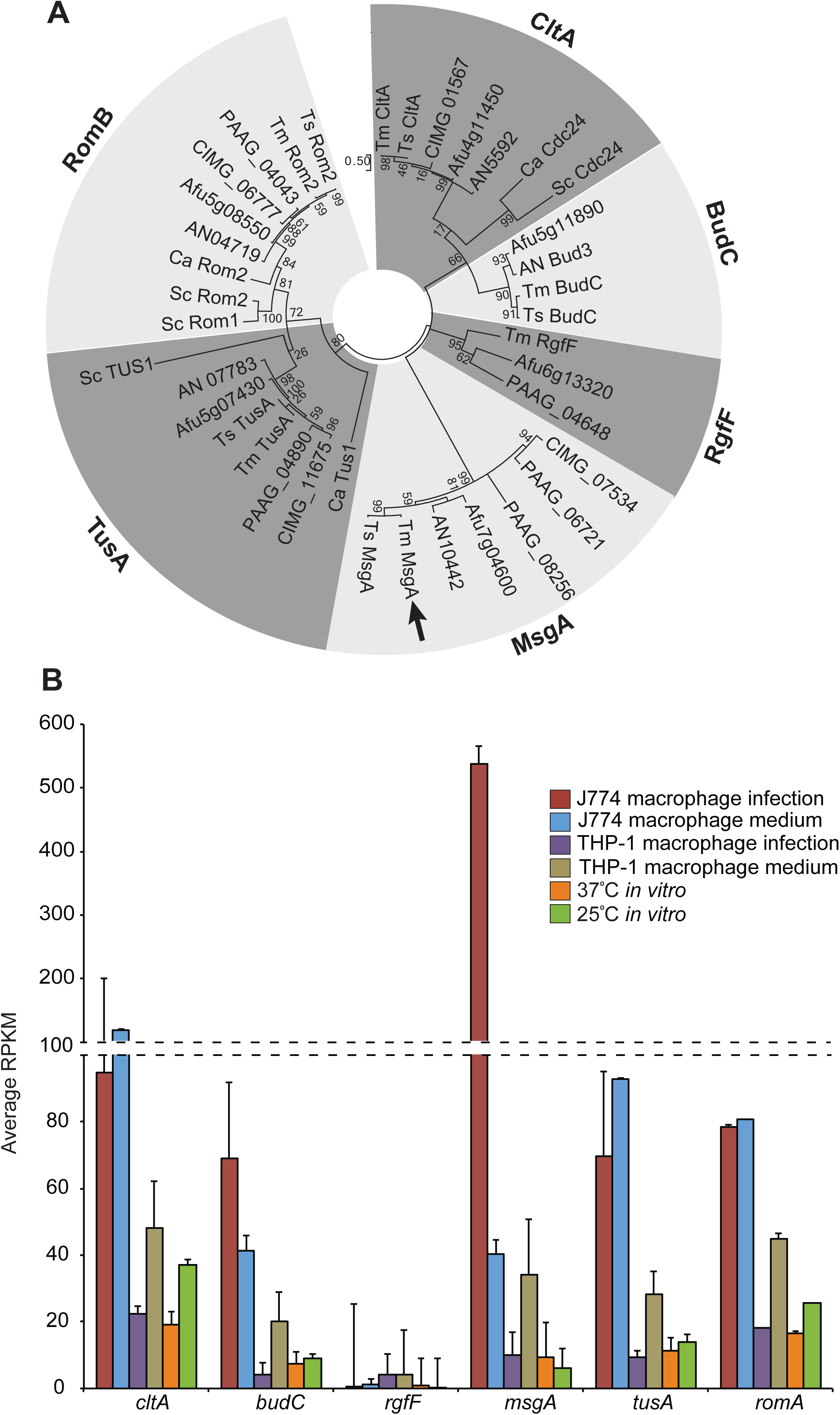
Relatedness of *T. marneffei* MsgA and other GEF-like proteins in fungi. (A) Bootstrapped relatedness tree of predicted Rho guanyl nucleotide exchange factor (GEF) proteins from a selection of dimorphic and monomorphic fungi. Black arrow indicates *T*. *marneffei* MsgA protein. The tree was generated using the maximum likelihood method based on JTT matrix model in CLUSTALW on MEGA7. Protein sequences are from *Saccharomyces cerevisiae* (Sc), *Candida albicans* (Ca), *Aspergillus nidulans* (AN), *Aspergillus fumigatu*s (Afu), *Histoplasma capsulatum* (HCBG), *Coccidioides immitis* (CIMG), *Paracoccidioides brasiliensis* (PAAG), *Talaromyces stipitatus* (Ts) and *Talaromyces marneffei* (Tm). A previously characterised, archetypal member denotes each clade of Rho GEFs where possible: CDC24-like (CtlA), Bud3-like (BudC), RgfF, MsgA, TUS1-like (TusA) and ROM1/ROM2-like (RomB). MsgA belongs to a distinct class of Rho guanyl nucleotide exchange factor (GEF) proteins. (B) Expression of *T*. *marneffei* Rho GEF encoding genes across several growth conditions. These conditions include 24 h growth inside J774 murine macrophages (J774 macrophage infection) and THP-1 human macrophages (THP-1 macrophage infection), 24 h growth in macrophage-free cell culture media (J774 macrophage medium and THP-1 macrophage medium), 4 days hyphal growth at 25°C(25°C*in vitro*) and yeast growth at 37°C(37°C*in vitro*) *in vitro*. RNA was extracted and quantified by RNAseq analysis. MsgA shows upregulated expression during growth inside J774 macrophages while the *ctlA*, *budC*, *tusA* and *romB* genes show expression across all conditions. The *rgfF* gene shows little to no expression under the tested conditions. Error bars represent the standard error of the mean.

Canonical RhoGEFs are composed of two distinct domains, a *<u>D</u>bl* <u>h</u>omology domain (DH) which catalyses the exchange of GDP for GTP within Rho GTPases and a <u>P</u>lekstrin <u>h</u>omology domain (PH) which assists in localisation of GEFs and regulation of their activity (reviewed in (Rossman et al. 2005)). Protein domain prediction analysis showed that *T*. *marneffei* RhoGEFs all contained a DH domain (Interpro ID: IPR000219) but showed differences in the composition of the second domain. CltA, TusA and RgfF have PH domains (Interpro ID: IPR001849), while RomA and BudC lack a second domain. However MsgA has a <u>BA</u>mphiphysin-<u>R</u>vs (BAR) homology domain (Interpro ID: IPR004148) downstream of the DH domain (Figure 1A), and no PH domain. BAR domains are known to interact with membranes and sense and promote membrane curvature as well as act as a binding platform for GTPases (Takei et al. 1999),(Habermann 2004). Additionally the predicted MsgA protein (1999 aa) is longer than other RhoGEFs, CtlA (932 aa), TusA (1781 aa), RomA (1217 aa), BudC (11546 aa) and RgfF (809 aa), and includes a large N-terminal region with no predicted domain motifs or localisation signals (Supplementary Figure 1A). This N-terminal sequence contains a 26 glutamic acid repeat (Supplementary Figure 1A). There is variation in the length of the acidic amino acid residue repeats in the MsgA orthologues from several clinical isolates of *T*. *marneffei*, ranging from 15 aa in isolates 3482 and 3841 to 26 aa in FRR 2161) (Supplementary Figure 1B). This is despite the high degree of sequence conservation outside this region. Additionally this repeat is either greatly reduced or absent in closely related non-pathogenic, pathogenic and dimorphic species (Supplementary Figure 1B and C). The significance of these dynamic acidic amino acids strings is unclear at this stage. Tandem repeated sequences (TRSs) have previously been shown to affect adhesion and host immune system evasion in other fungi (Verstrepen et al. 2005; Verstrepen et al. 2004). A recent study of the genome structure of *T*. *marneffei* identified an increase in number of genes with TRSs compared to three other pathogenic and non-pathogenic filamentous fungi (Yang et al. 2013).

To examine the expression of the six genes encoding RhoGEF-like proteins in *T*. *marneffei*, including *msgA,* we queried the RNAseq analysis data (Weerasinghe *et al* in prep), which included gene expression profiles during *in vitro* hyphal growth at 25°C, *in vitro* yeast growth at 37°C, intracellular macrophage growth in J774 murine macrophages and THP-1 human macrophages, as well as growth in macrophage-free cell culture media. Most GEFs showed minimal (*ctlA*, *tusA budC* and *romA*) to no (*rgfF*) expression during yeast growth, either *in vitro* or during macrophage infection (Figure 1B). In contrast *msgA* showed upregulated expression specifically in the J774 macrophage infection condition, with a 13.4 fold increase compared to the J774 *in media* condition. This data suggested that *msgA* might have a unique role during intracellular growth.

### MsgA is required for the formation of yeast inside host cells

To characterise the role played by *msgA* during morphogenesis of *T*. *marneffei*, the gene was cloned, a deletion construct generated and this was used to create a deletion strain (*ΔmsgA*) by DNA-mediated transformation. To confirm that any resulting phenotypes were caused by this gene deletion, the *ΔmsgA* strain was complemented with the wildtype (*msgA*^*+*^) allele targeted to the *niaD* locus. The wildtype (*msgA*^*+*^), Δ*msgA* and Δ*msgA msgA*^*+*^ strains were grown in liquid BHI medium at 25°C(hyphal) and 37°C(yeast) *in vitro* for 4 and 6 days respectively. At 25°Cconidia from all three strains germinated normally and exhibited wildtype morphology, growing as polarized, branched septate hyphae. Similarly, at 37°Cconidia from all three strains germinated and subsequently produced yeast cells (Figure 2A). Thus the Δ*msgA* and Δ*msgA msgA*^*+*^ strains were indistinguishable from wildtype under *in vitro* growth conditions. This is consistent with the expression data in which the *msgA* transcript is present at very low levels during *in vitro* growth (Figure 1B).

**Figure 2.**
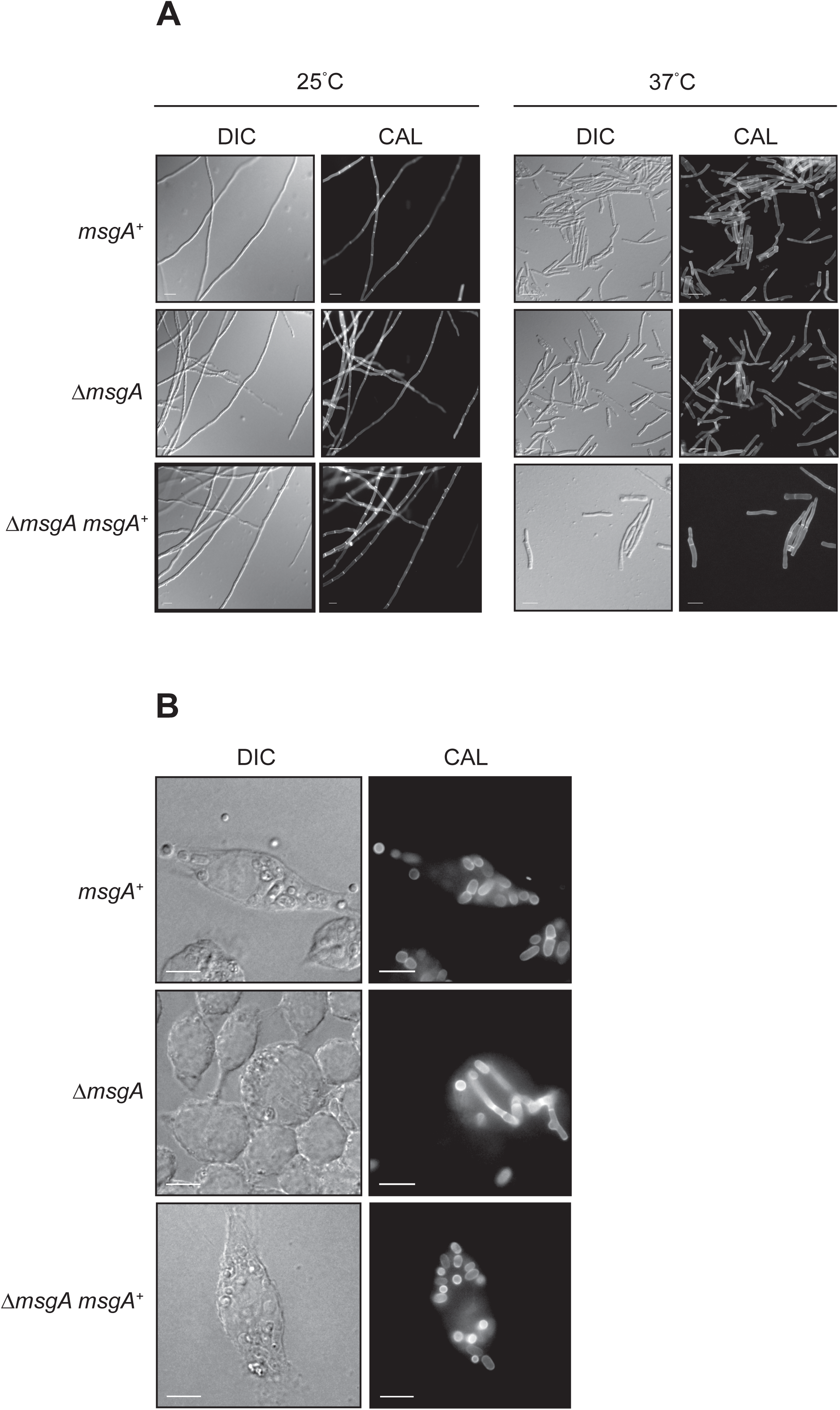
Deletion of *msgA* leads to aberrant growth inside macrophages. (A) The wildtype and Δ*msgA* strains were grown on BHI medium at 25°Cfor 5 days and 37°Cfor 6 days, stained with calcofluor (CAL) and examined. Both strains show indistinguishable growth characteristics with respect to growth rate and morphology in the hyphal (25°C) and yeast (37°C) form. (B) LPS activated J774 murine macrophages were infected with conidia from the wildtype, Δ*msgA* and complemented Δ*msgA msgA*^*+*^ strains and examined microscopically. At 24 h post infection wildtype *T*. *marneffei* within J774 macrophages produced numerous ovoid yeast cells that divided by fission. In contrast the Δ*msgA* mutant produced aberrantly shaped filaments, in addition to yeast cells. The complemented strain (Δ*msgA msgA*^*+*^) was indistinguishable from the wildtype. (C) The effects of deleting *msgA* on morphogenesis in macrophages were quantified at the 24 h post infection time point. The Δ*msgA* mutant strain showed 37.4±0.49% septate filaments compared to the wildtype and complemented Δ*msgA msgA*^*+*^ strains, which had 0.6±0.02% and 2.5±0.6% respectively. Additionally, the Δ*msgA* strain showed a reduction in the number of yeast cells during macrophage infection with 55.8±1.85% compared to 81.8±1.89% and 86±0.56% in the wildtype and Δ*msgA msgA*^*+*^ strains respectively. Error bars represent the standard error of the mean with t-test values falling in the following range ** ≤0.05. Images were captured using differential interference contrast (DIC) or with epifluorescence to observe calcofluor stained fungal cell walls (CAL). Scale bars are 10 µm.

When these stains were used to infect LPS activated J774 macrophages the wildtype and Δ*msgA msgA*^*+*^ strains were phagocytosed and germinated to produce small ellipsoid yeast cells that divided by fission after 24 h. In contrast macrophages infected with Δ*msgA* conidia were equally phagocytosed but contained more branched, septate filamentous-like cells after 24 h (Figure 2B). Counts of the number of *T*. *marneffei* cells with at least one septum across approximately 100 infected macrophages showed that the wildtype and Δ*msgA msgA*^*+*^ strains had 0.6±0.02% and 2.5±0.6% septate cells, respectively, while the Δ*msgA* strain had 37.4±0.49% septate cells. Concomitantly, the Δ*msgA* strain showed a reduction in the number of yeast cells during macrophage infection with 55.8±1.85% compared to 81.8±1.89% and 86±0.56% in the wildtype and Δ*msgA msgA*^*+*^ strains respectively. Longitudinal cell length measurements showed that yeast cells produced by the Δ*msgA* strain were on average 1.4 times longer than both the wildtype and the Δ*msgA msgA*^*+*^ strains. This shows that *msgA* is important for correct yeast cell morphogenesis exclusively during macrophage infection.

Yeast cell morphogenesis *in vitro* occurs over days, rather than hours as seen inside host cells. So an explanation for the aberrant yeast cell morphology inside macrophages is that the cells are growing more slowly in the mutant. To test this hypothesis conidia of the wildtype, Δ*msgA* and Δ*msgA msgA*^*+*^ strains were used to infect J774 macrophages and examined after extended incubation for 48 h post infection. At this time point macrophages infected with all three strains contain a large number of yeast cells dividing by fission. Macrophages infected with the wildtype or Δ*msgA msgA*^*+*^ strains showed similar yeast cell morphology that was short and oval shaped. In contrast, macrophages infected with the Δ*msgA* strain contained long and cylindrical yeast cells with at least one septum compared to the long multi septate, filaments observed at 24 h post infection (Figure 3A). The Δ*msgA* strain yeast cells were 2.3 times longer than wildtype,being 8.9±0.57μm, compared to wildtype and Δ*msgA msgA*^*+*^ that were 3.9±0.12μm and 4.1±0.50μm respectively (Figure 3B). As the Δ*msgA* strain does not show germination defects either *in vitro* or during macrophage infection when compared to wildtype (Supplementary Table 3) and there are no morphological differences between the wildtype and Δ*msgA* strains at 4, 6 or 8 h post-infection *ex vivo* (data not shown), the elongated yeast cell morphology observed in the Δ*msgA* strain during macrophage infection is unlikely to reflect a delay in germination. Rather it points to a requirement of *msgA* in maintaining the compact, ellipsoid yeast cell morphology specific to macrophage infection.

**Figure 3.**
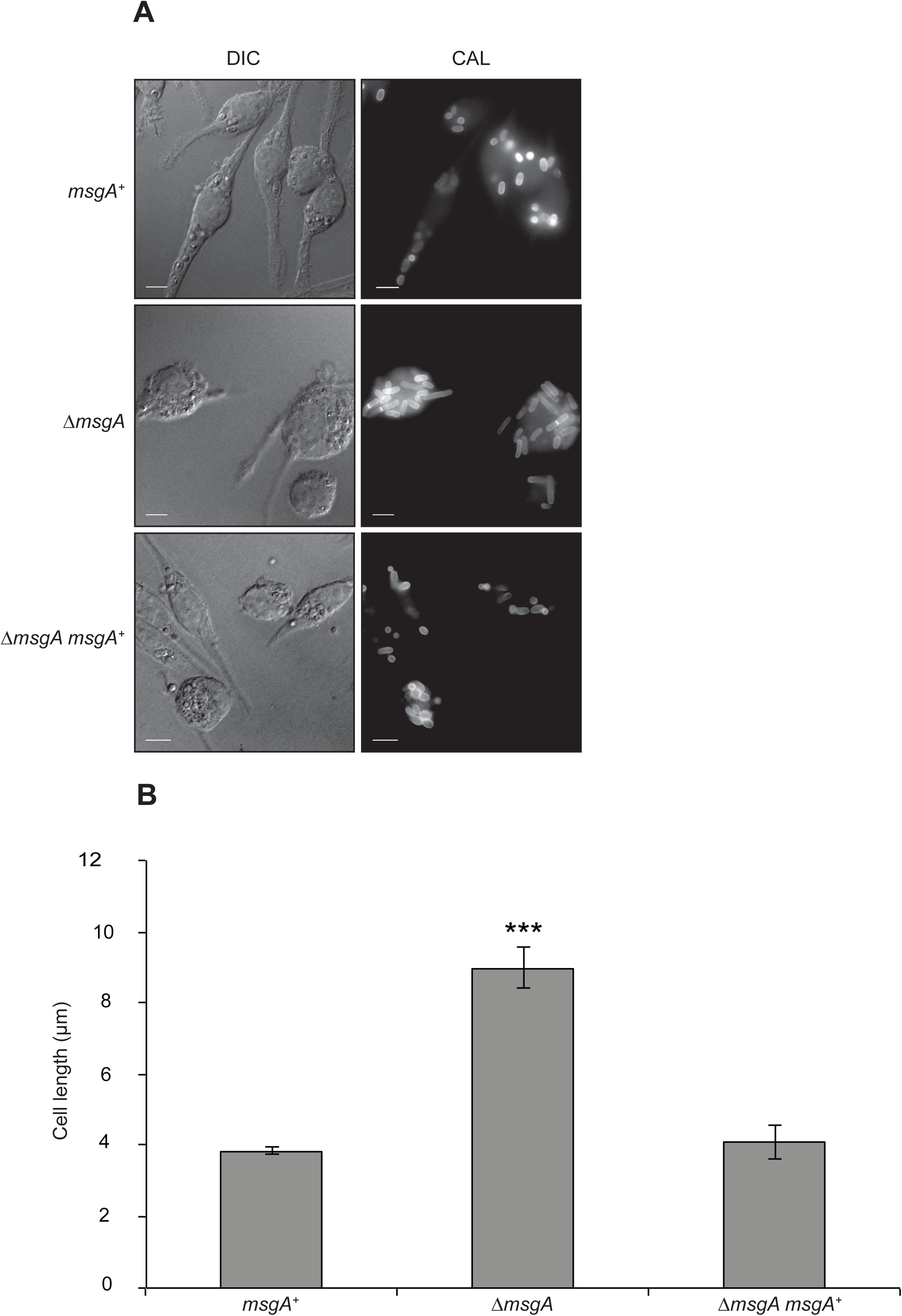
Yeast cell formation defects in the ΔmsgA strain are not resolved with prolonged incubation in macrophages. LPS activated J774 murine macrophages were infected with conidia from the wildtype, Δ*msgA* and complemented Δ*msgA msgA*^*+*^ strains and examined microscopically after prolonged incubation. (A) At 48 h post infection wildtype (*msgA*^*+*^) *T*. *marneffei* within J774 macrophages retain their ovoid morphology producing numerous yeast cells that divided by fission. However the septate filaments produced by the Δ*msgA* mutant strain at 24h post infection continued to grow, showing increased length, rather than breaking down to form short ellipsoid yeast cells (B) The effect of deleting *msgA* on morphogenesis in macrophages was quantified at the 48 h post infection time point. Wildtype yeast cells showed an average length of 3.9±0.12μm compared to the Δ*msgA* mutant strain, which produced yeast cells of an average cells length of 8.9±0.57μm, approximately 2.3 times longer than wildtype. The complemented strain (Δ*msgA msgA*^*+*^) produced yeast cells of 4.1±0.50μm in length and was comparable to wildtype. Error bars represent standard error of the mean with t-test values falling in the following range *** ≤0.001.

### Aberrant morphogenesis of the msgA mutant is not due to defective host cell sensing

The germination and morphogenesis of dormant conidia, which are produced during the hyphal growth phase at 25°C, into yeast cells differs *in vitro* compared to inside host cells. *In vitro* the conidia produce a polarised growth tip and generate multinucleate, multicellular hyphae before undergoing coupled nuclear and cell division to produce arthroconidiating hyphae. Fragmentation of these hyphae at septal junctions liberates uninucleate yeast cells. Inside host cells conidia undergo isotropic growth upon germination and form yeast cells directly. The morphology of Δ*msgA* mutant cells in J774 macrophages resembled arthroconidiating hyphae, so one explanation for the mutant phenotype is that they can no longer accurately sense the host environment. To test this the expression of four genes known to be expressed at 37°C*in vitro*, but with little to no expression during J774 macrophage infection was examined (Weerasinghe *et al*., in prep). RNA was extracted from the wildtype and Δ*msgA* strains grown at 37°C*in vitro* or during J774 macrophage infection and used for RT-PCR analysis. While the four genes showed a clear transcript in the wildtype grown at 37°C*in vitro*, no expression was observed during the J774 macrophage infection condition for either wildtype or Δ*msgA* (Supplementary Figure 2). This suggests that the filaments and elongated yeast cells seen in the Δ*msgA* strain are not similar to arthroconidial hyphae and yeast cells produced during wildtype growth *in vitro* at 37°C, and *msgA* has a specific role in maintaining cell shape during macrophage infection.

### MsgA also plays a role in the formation of yeast cells inside human cells

The expression level of *msgA* in human THP-1 macrophages is very low compared to murine J774 cells at the 24h time point. To determine if the aberrant yeast cell morphogenesis phenotype of the Δ*msgA* strain was limited to growth in J774 macrophages, THP-1 cells were infected with conidia from wildtype, Δ*msgA* and Δ*msgA msgA*^*+*^ strains and observed at 24 h post infection (Supplementary Figure 3A). The yeast cells displayed an increase in length (8.6±0.25μm) compared to wildtype (4.9±0.09μm) but were not as long, relative to wildtype, when growing inside J774 mouse cells (Supplementary Figure 3B). This suggests that *msgA* also plays a role in yeast cell morphogenesis inside THP- 1 macrophages, albeit less pronounced.

### Overexpression of msgA produces aberrant hyphal morphology during growth at 25°C

The *msgA* gene is important for yeast cell morphogenesis in macrophages and shows very little expression at 25°Cand 37°C*in vitro* (Figure 1A). In an attempt to gain more insight into the cellular activity of MsgA a construct that contained *msgA* under the control of a xylose inducible promoter, *xylP*, was generated and used to transform *T*. *marneffei* strain G809 (Materials and methods). The wildtype and *xylP(p)*::*msgA*, strains were grown in BHI medium with (inducing) or without (non-inducing) 1% xylose (Materials and Methods). On non-inducing medium the *xylP(p)::msgA* strain was indistinguishable from wildtype at both the macroscopic and microscopic levels. On inducing medium the *xylP(p)::msgA* strain produced slightly swollen hyphae that displayed increased septation and hyper-branching along the hyphal length (Figure 4A). The distance between adjacent septa was 38.6±0.9 μm for the wildtype, 36.3±1.1 μm for the Δ*msgA* strain and 6.6±0.18 μm for the *xylP(p)::msgA* strain. The frequency of branching in sub-apical cells along hyphae was 1.15±0.11 branches per 100 μm of hyphal length for the wildtype, 1.06±0.51 for the Δ*msgA* strain and 11.05±0.81 for the *xylP(p)::msgA* strain. Branching in apical cells was not observed in any of the strains and there was no significant difference in the nuclear index between these strains (data not shown). Extended incubation up to 6 days under inducing conditions did not lead to the septate hyphae breaking down to form yeast cells. Therefore, the data shows that MsgA activity is required for specifying cell shape in host cells and can drive cell shape changes *in vitro*, possibly by affecting cell division.

**Figure 4.**
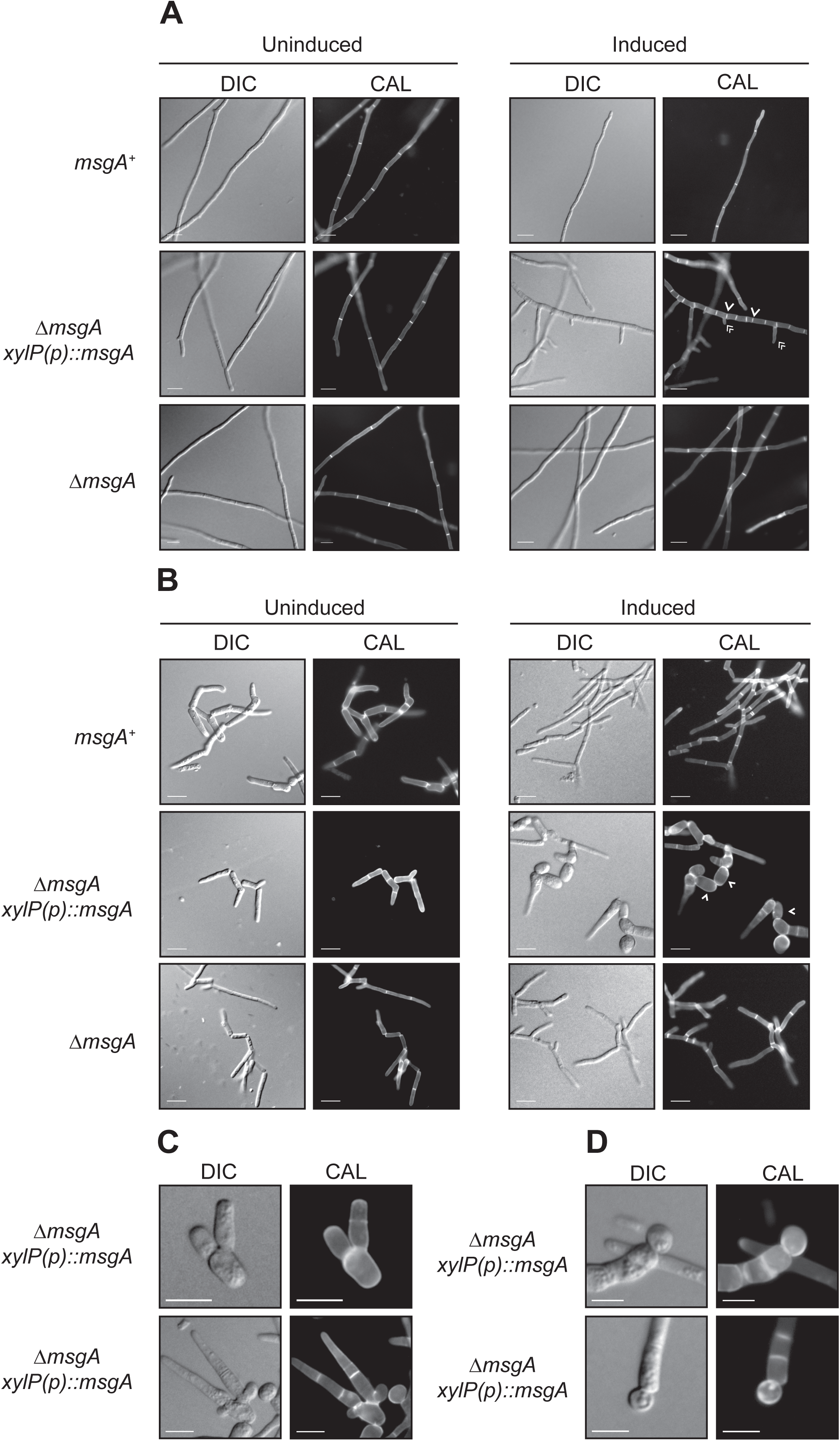
Induced overexpression of *msgA* produces aberrant hyphae at 25°Cand yeast cells with intracellular morphology at 37°C*in vitro*. The wildtype (*msgA*^*+*^), inducible (Δ*msgA xylP(p)*::*msgA*) and deletion mutant (Δ*msgA*) allele strains were grown in liquid BHI medium supplemented with either 1% glucose (Uninduced) or 1% xylose (Induced) for 5 days at 25°C(A) or 6 days 37°C(B). Under uninduced conditions the Δ*msgA xylP(p)*::*msgA* strain is indistinguishable from Δ*msgA* at both temperatures. (A) On inducing medium at 25°Cthe Δ*msgA xylP(p)*::*msgA* strain shows increased septation (single arrowheads) and branching (double arrowheads) along the entire length of the hyphae but all other morphological and growth characters were indistinguishable amongst the strains. (B) On inducing medium at 37°C the Δ*msgA xylP(p)*::*msgA* strain produced yeast cells that are rounder and greatly reduced in length compared to the Δ*msgA* (single arrowheads). These yeast cells were dividing by fission and resembled yeast cells produced during growth inside macrophages. (C) Continuous induction over six days of growth of the Δ*msgA xylP(p)*::*msgA* strain produces yeast cells that have polarity defects at division and (D) a proportion of yeast cells appear to be dividing by budding (white triangle). Images were captured using differential interference contrast (DIC) or with epifluorescence to observe calcofluor stained fungal cell walls (CAL). Scale bars are 10 µm.

### Overexpression of msgA at 37°Cin vitro mimics yeast cell morphogenesis during macrophage infection

Loss of *msgA* leads to the production of long and cylindrical yeast cells with *in vitro* morphology during macrophage infection rather than the wildtype short, ellipsoid cells. To test if MsgA could drive the development of the intracellular macrophage growth yeast cell morphology at 37°C*in vitro*, yeast cells from the wildtype and *xylP(p)::msgA*, strains were transferred to BHI medium at 37°Cboth with (inducing) and without (non-inducing) 1% xylose (Materials and Methods). Under non-inducing conditions, the *xylP(p)::msgA* strain was indistinguishable from wildtype, whereas on inducing medium the *xylP(p)::msgA* strain produced yeast cells that were rounder and greatly reduced in length compared to wildtype (Figure 4B), resembling those produced by *T*. *marneffei* during J774 macrophage infection (Figure 4B). These yeast cells also displayed patchy, uneven chitin staining. Additionally the arthroconidial filaments that produce these yeast cells display aberrant chitin deposition and an increased septation frequency immediately adjacent to the fission division sites (similar to the induced *xylP(p)::msgA* strain at 25°C).

I*n vitro* generated yeast cells consist of a mixture of arthroconidia (single cells formed by the separation of hyphal cells) and *bona fide* yeast cells (arthroconidial cells that have divided at least once). To determine if MsgA could drive the development of yeast cells directly from conidia at 37°C*in vitro*, as happens inside host cells, conidia from the wildtype, Δ*msgA* and *xylP(p)::msgA* strains were grown under continuous induction in BHI medium with 1% xylose. After 6 days the wildtype and Δ*msgA* strains produce elongated yeast cells under both inducing and non-inducing conditions. In contrast, the *xylP(p):::msgA* strain produced small ellipsoid yeast cells similar to those observed when *T*. *marneffei* is growing inside host cells (Supplementary Figure 4). These yeast cells appear to be derived from swollen arthoroconidial filaments rather than directly from conidia. Additionally, wildtype yeast cell formation occurs by fission driven separation from either pole of the primary yeast cell (Supplementary Figure 4). However, under continuous induction the *xylP(p)::msgA* strain produces a proportion of aberrantly shaped yeast cells, with uneven chitin deposition, that appear to produce yeast cells from multiple fission sites at the poles, often resulting in the production of two yeast cells from a singe pole (Figure 4C). Sometimes these aberrant yeast cells, as well as long filamentous cells, produce cells that have lost their ellipsoid shape and seem to be dividing by budding rather than fission (Figure 4D). Thus while the overexpression of *msgA in vitro* is able to recapitulate the phenotype of *T*. *marneffei* yeast cells grown within macrophages, continuous overexpression impedes yeast cell growth polarisation and results in inappropriate division.

### The MsgA protein is involved in conidial production during asexual development

During asexual development at 25°C, wildtype *T*. *marneffei* colonies are composed of vegetative hyphae that produce asexual differentiated structures (conidiophores) from which green-pigmented asexual spores (conidia) are generated by basipetal budding (Borneman et al. 2000). The Δ*msgA* mutant strain produced fewer conidia than wildtype, resulting in a paler colonial appearance (Supplementary Figure 5A). Conidial counts of wildtype, Δ*msgA* and Δ*msgA msgA*^*+*^ strains revealed that the Δ*msgA* strain produced 5.3×10^8^±0.4 conidia/ml while the wildtype produced 23.9×10^8^±0.9 conidia/ml. The complemented Δ*msgA msgA+* strain did not completely rescue the phenotype of the Δ*msgA* mutant strain with conidiation levels at 15.9×10^8^±1.1 conidia/ml (Supplementary Figure 5B). The complementation allele included 891 bp of promoter sequence, however examination of the 3624 bp region between the *msgA* start codon and the upstream gene (PMAA_089490) identified four putative recognition sites for asexual development transcriptional regulators BrlA (5’-MRAGGGT-3’, at −1162 and –3067) and AbaA (5’-CATTCY-3’ at -339 and -1711). Of these only one (AbaA at -339) site was within the region included in the reintroduced allele. This may explain the partial complementation observed in the Δ*msgA msgA*+ strain. A similar phenotype has been observed for the *drkA* gene in *T*. *marneffei*, which was attributed to the absence of BrlA and AbaA recognition sites (Boyce et al. 2011).

To test this hypothesis conidial density of the *msgA* over expression strain (*xylP(p)::msgA*) was measured with and without induction. In the absence of induction the number of conidia produced the *xylP(p)::msgA* strain was comparable to the Δ*msgA* mutant strain, being 7.7×10^8^±0.8 conidia/ml. However in the presence of 0.5% xylose induction the conidiation level in the *xylP(p)::msgA* strain was 35.2×10^8^±1.5 conidia/ml, which is comparable to wildtype which produced 33×10^8^±1.5 conidia/ml (Supplementary Figure 5B). This suggests that promoter elements excluded in the *msgA*^*+*^ complementation allele are necessary for the proper expression of *msgA* during asexual development, and highlights a role for *msgA* during conidial production.

### MsgA shows cell membrane localisation during macrophage growth

To investigate the localization of MsgA, the mCherry coding sequence was inserted into the C-terminal end of MsgA, after the DH domain. The *msgA::m*Cherry fusion construct was targeted to the *niaD* locus in the *ΔmsgA* (G1003) strain. This strain was used to infect J774 murine macrophages and wildtype yeast cell morphology was evident after 24 h showing that the fusion allele complemented the Δ*msgA* phenotype during macrophage infection (Figure 5A). Immunostaining of these infected cells with an anti-mCherry antibody and calcofluor co-staining showed that MsgA-mCherry specifically localized to yeast cells but not to ungerminated conidia (Figure 5B). Localization was observed around the cell periphery and in distinct punctate vesicular structures within the cytoplasm. Colocalization with calcofluor stained cell walls and septa was not observed, either at nascent septation sites prior to, or immediately after, cell wall deposition (Figure 5B-C). However localization of MsgA was observed in the region immediately adjacent to the cell wall during cell separation (Figure 5D-E). Brighter punctate structures of MsgA localization can be seen at the sites of cytokinesis in cells that are actively undergoing fission division. Thus MsgA is localised to the cell membrane and vesicles during macrophage infection, with an increased presence at cell division sites late in the cell division process.

**Figure 5:**
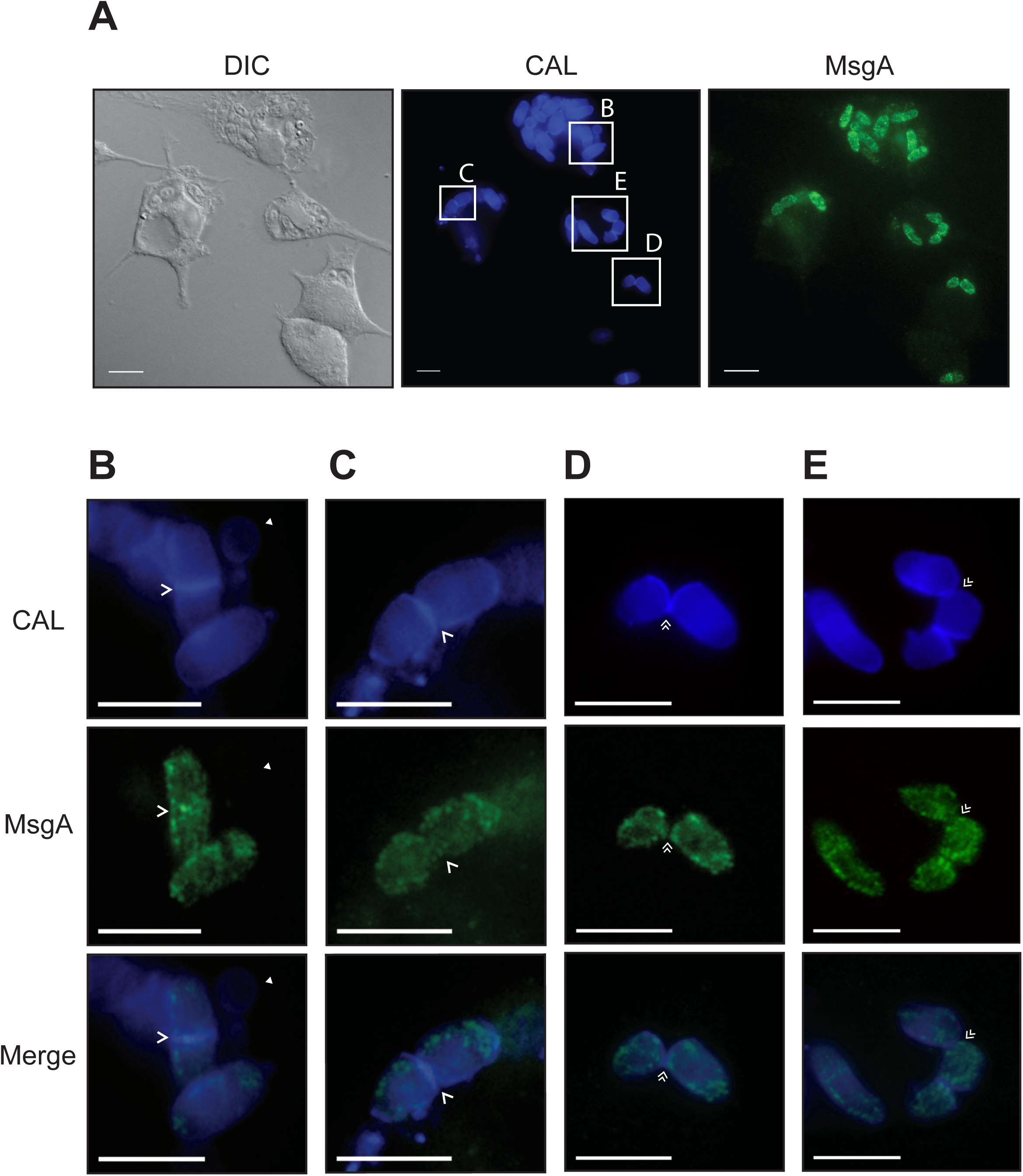
MsgA localisation during growth inside J774 murine macrophages. LPS activated J774 murine macrophages were infected with a strain expressing the *msgA*::mCherry fusion gene and incubated for 24 h. Cells were fixed and the MsgA-mCherry fusion (MsgA) detected using an anti-mCherry rat monoclonal (3F10) primary and an anti-rat ALEXA488 goat secondary antibody. Yeast cells were also stained with calcofluor (CAL) to highlight the fungal cell wall. The MsgA and CAL panels were also merged (Merge) to assess relative localisation (A). MsgA was only evident in actively growing yeast cells and not in ungerminated conidia (solid arrowhead) (B). In yeast cells MsgA is localised to the cell membrane, immediately adjacent to the cell wall, and shows punctate localisation in the cell membrane periphery, as well as both within the cytoplasm (D-E). Localization is not evident either at nascent septation sites prior to (B), or immediately after (C)(single arrowheads), cell wall deposition. Instead it appears in a region immediately adjacent to septation sites during cell separation (D-E) (double arrowheads). Images were captured using differential interference contrast (DIC) or with epifluorescence to observe calcofluor and ALEXA488 stained fungal features. Scale bars are 10 µm.

### The BAR domain of MsgA is essential for proper yeast morphogenesis during macrophage infection

In order to examine the role of the conserved domains of MsgA in the phenotypes observed during macrophage infection, mutant alleles were generated which deleted the DH (*msgA*^ΔDH^) or BAR domains (*msgA*^ΔBAR^). Domain deletion constructs were targeted to the *niaD* locus of the Δ*msgA* (G1003) strain. These mutant strains were compared to the wildtype, the original deletion strain (Δ*msgA*) and the wildtype complementation strain (Δ*msgA msgA*^*+*^). Both the *msgA*^ΔDH^ and *msgA*^ΔBAR^ strains were indistinguishable from the control strains during *in vitro* growth at 25°Cand 37°C(data not shown). When conidia from these strains were used to infect LPS activated J774 murine macrophages numerous yeast cells that were dividing by fission were present for the wildtype, Δ*msgA msgA*^*+*^ and *msgA*^ΔDH^ strains after 24 h. In contrast macrophages infected with Δ*msgA* and *msgA*^ΔBAR^ strains contained both septate yeast as well as long septate filament cells (Figure 6A). Macrophages infected with the *msgA*^ΔBAR^ strain contained 39.1±0.6% filaments of the total fungal load when compared to the *msgA*^ΔDH^ strain, which contained 0.8±0.4% filaments (Figure 6B). Similar to the Δ*msgA* strain, the *msgA*^ΔBAR^ strain contained fewer yeast cells compared to the *msgA*^ΔDH^ and wildtype strains (Figure 6B). Here the *msgA*^ΔBAR^ strain produced 59.1±2.7% yeast cells compared to 84.4±0.5% formation rate in the *msgA*^ΔDH^ strain. Thus the *msgA*^ΔBAR^ allele did not complement the Δ*msgA* mutation, indicating that the BAR domain is essential for normal yeast morphogenesis during macrophage infection. Both domain deletion alleles were also introduced into the wildtype strain but did not alter the wildtype phenotype under either the *in vitro* or intracellular macrophage growth conditions tested, suggesting that these alleles do not have a dominant interfering effect on the function of MsgA (data not shown).

**Figure 6:**
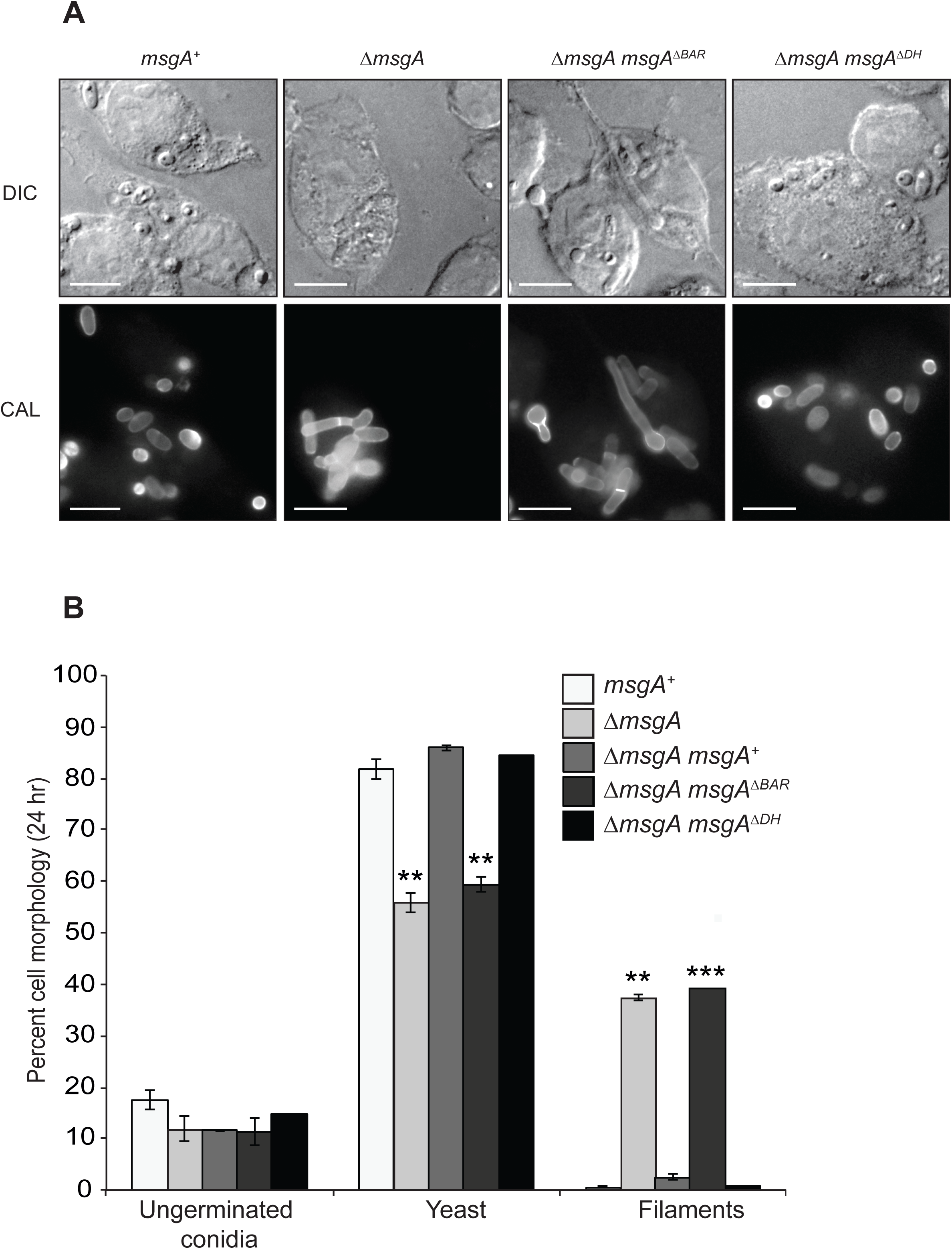
The BAR domain of *msgA* contributes to morphology during growth inside macrophages. (A) LPS activated murine macrophages infected with conidia of either the wildtype (*msgA*^*+*^), Δ*msgA*, 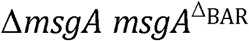 or 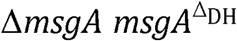 strain. After 24 h cells were fixed, stained with calcofluor (CAL) and examined microscopically. Numerous small ellipsoid yeast cells dividing by fission were observed in macrophages infected with the wildtype and 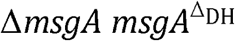 strains. In contrast macrophages infected with Δ*msgA* and 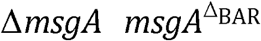 strains contained septate filaments and elongated yeast-like cells. (B). The effects of the *msgA* alleles on morphogenesis in macrophages were quantified at the 24 h post infection time point. The 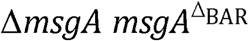 mutant strain shows 39.1±0.6% more filaments and 22.7%±0.5 fewer yeast cells compared to wildtype. These numbers are equivalent to those for the Δ*msgA* strain. In contrast the 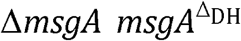 strain was comparable to the wildtype and complemented Δ*msgA* strain. Error bars represented SEM with t-test values falling in the following range ** ≤0.05 and *** ≤0.001. Images were captured using differential interference contrast (DIC) or with epifluorescence to observe calcofluor stained fungal cell walls (CAL). Scale bars are 10μm.

To assess whether the DH or BAR deletion mutations affect the localization of MsgA during macrophage infection, mCherry tagged *msgA*^ΔDH^ and *msgA*^ΔBAR^ constructs were generated and targeted to the *niaD* locus in the Δ*msgA* (G1003) strain. These strains were used to infect J774 murine macrophages and, as observed for the non-mCherry fusion alleles, wildtype yeast cell morphology was evident after 24 h for the *msgA*^ΔDH^::mCherry allele but not for the *msgA*^ΔBAR^::mCherry allele (Figure 7). Immunostaining of these infected cells with an anti-mCherry antibody and calcofluor co-staining showed that the localisation pattern for the MsgA^ΔDH^-mCherry was the same as that of the MsgA-mCherry with punctate localisation in the cytoplasm and at the cell membrane and none at the cell wall either during cell wall deposition at the septa or during cytokinesis. In contrast, the MsgA^ΔBAR^-mCherry gene product showed mislocalisation to the cell wall where it co-staining with calcofluor. The MsgA^ΔBAR^-mCherry gene product did not show the punctate pattern of localisation within the cytoplasm like wildtype and was not localized to nascent septation sites prior to or immediately after cell wall deposition (Figure 7). However MsgA^ΔBAR^-mCherry showed weak localisation at the cell wall of septation sites during cell separation (Figure 7). This suggests the possible late recruitment of MsgA to the cell separation complex.

**Figure 7:**
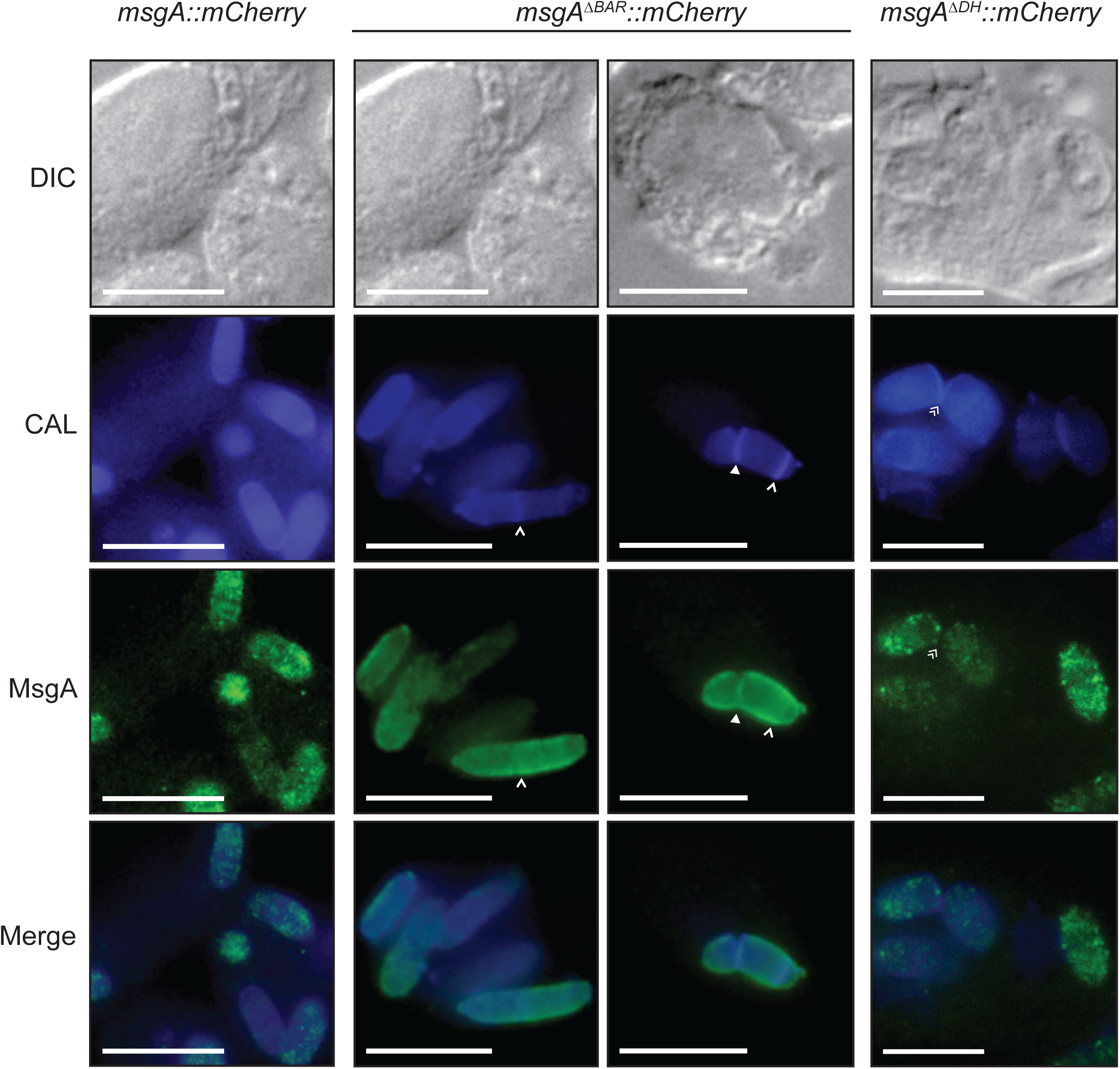
The BAR domain is necessary for the localisation of MsgA during growth inside macrophages. LPS activated murine macrophages infected with conidia from the wildtype (*msgA::mCherry*), 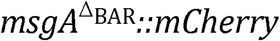 or 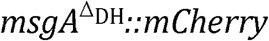 strain. After 24 h cells were fixed and the MsgA-mCherry fusion (MsgA) alleles was detected using an anti-mCherry rat monoclonal (3F10) primary and an anti-rat ALEXA488 goat secondary antibody. Yeast cells were also stained with calcofluor (CAL) to highlight the fungal cell wall. Both MsgA and CAL panels were overlayed to indicate overlapping localisation (Merge). The wildtype *msgA::mCherry* gene product showed clear punctate cell membrane localisation with puncta also within the cytoplasm (*msgA::mCherry* panel). This pattern of localisation was also evident for the *msgA*^*ΔDH*^*::mCherry* gene product (*msgA*^*ΔDH*^*::mCherry* panel). However the *msgA*^*ΔBAR*^*::mCherry* gene product has lost its punctate pattern of localisation and appears uniformly within the cytoplasm. Additionally the *msgA*^*ΔBAR*^*::mCherry* gene product was mislocalized to the cell wall co-staining with calcofluor throughout the cell periphery (*msgA*^*ΔBAR*^*::mCherry* panel). As with the wildtype *msgA::mCherry* strain (Figure 5) the *msgA*^*ΔDH*^*::mCherry* gene product localized to the region immediately adjacent division septum during cell separation (double arrowheads). In contrast the *msgA*^*ΔBAR*^*::mCherry* gene product was mislocalized to the cell wall, co-staining with calcofluor, during cell separation (solid arrowhead) but not at nascent septation sites prior to (single arrowheads) cell wall deposition. Images were captured using differential interference contrast (DIC) or with epifluorescence to observe calcofluor (CAL) and ALEXA488 fluorophores. Scale bars are 10 µm.

## Discussion

Survival and proliferation of microbial pathogens in a host relies on their ability to cope with host defence mechanisms and acquire nutrients for growth. For a number of prokaryotic and eukaryotic microbial pathogens this is coupled with the ability to grow inside cells of the host. In intracellular dimorphic pathogenic fungi, the ability to switch from a multicellular hyphal growth form into a unicellular yeast form that is more spatially suited to residing within host cells, is crucial for pathogenicity (Nemecek et al. 2006; Webster & Sil 2008; Beyhan et al. 2013). An important inducer of this switch is a shift to 37°C(mammalian body temperature), and this coincides with the conversion of infectious propagules to a pathogenic form. This study examined a previously uncharacterised *Dbl* homology/BAR domain protein encoded by *msgA*, which is strongly upregulated in the dimorphic, human-pathogenic fungus *T*. *marneffei* during J774 murine macrophage infection and is critical for yeast morphogenesis in macrophages. It was demonstrated that *msgA* does not play a role *in vitro*, during yeast or hyphal cell morphogenesis at either 37°Cor 25°C, but does affect asexual development. Factors that are important for both yeast cell morphogenesis and asexual development have been identified previously. In particular studies of the cell signalling pathways in *T*. *marneffei* have uncovered similar effects for PakB, the *CLA4* homologue encoding a p21-activated kinase, in yeast cell formation particularly during macrophage growth (Boyce & Andrianopoulos 2007). Together the data indicate that morphogenesis in *T*. *marneffei* during infection may respond to a host internalisation triggered mechanism that induces the expression of *msgA* and other associated genes, and that the activation of this pathway drives cellular processes that responds to the host environment.

### MsgA is involved in intracellular morphogenesis

The *msgA* gene has a unique expression profile that foreshadows its contribution to yeast cell morphogenesis during macrophage infection. In comparison to the other six *Rho* GEF encoding genes, *msgA* shows high level and specific expression in *T*. *marneffei* during J774 murine macrophage infection. Consistent with this expression pattern the *msgA* deletion strain displayed septate and branched filament production during intracellular growth but wildtype yeast cell morphogenesis *in vitro* at 37°C. These aberrant *ex vivo* generated filaments were morphologically more similar to arthroconidia/yeast cells produced at 37°C*in vitro* but they failed to show expression of a number of genes known to be expressed exclusively in *in vitro* generated yeast cells, suggesting that the defect is unlikely to be the result of a failure to detect the local environment. A similar, albeit less severe, phenotype was observed when THP-1 human macrophages were used. An alternative explanation for the morphogenesis defect is that the phagocytosed conidia germinate aberrantly, however kinetic studies of germination for the *msgA* deletion strain failed to identify any differences compared to the wildtype under any condition. Coupled with the observation that overexpression of *msgA* at 37°C*in vitro* drives the formation of shorter ellipsoid yeast cells that resemble yeast during intracellular macrophage growth suggests that *msgA* is a key determinant of cell shape and identity and is sufficient to redirect cellular morphogenetic programs towards infection-specific yeast cell formation.

Unlike other dimorphic pathogens where the yeast phase is marked by budding division, *T*. *marneffei* yeast cells divide by fission, so yeast and hyphal cells share the polarised growth characteristics, up to the point of cell division. At cell division hyphal cells lay down a cell wall to generate a new cell compartment with no associated cell separation, whilst yeast cells undergo cell separation. The formation of elongated, filament-like yeast cells that are sometimes multiseptate by the Δ*msgA* mutant strain during infection points to a disruption in the coordination of these processes and suggests that MsgA plays a vital role in processes that involve cell division (septation) and cell separation. While its precise role remains to be determined, the redistribution of MsgA from the entire cell membrane to that adjacent to the newly formed septum and before cell separation suggests that it is involved more at the cell separation stage than septation. The smaller, ellipsoid yeast cells produced by overexpressing *msgA* at 37°C*in vitro* and the defect in conidiogenesis for the Δ*msgA* mutant strain are both consistent with this hypothesis that *msgA* is important for cell division and separation. Additonally, overexpressing *msgA* at 25°C*in vitro* results in increased septation along the hyphae producing shorter cellular compartments. Unlike yeast cell growth these compartments do not separate into individual cells, which is probably due to the absence of expression and/or activity of the yeast-specific machinery necessary for separation.

It is clear that *msgA* plays a fundamental role in the morphogenesis of yeast cells growing inside host cells and for conidial production during asexual development, The *pakB* gene of *T*. *marneffei*, encoding a p21-activated kinase, also plays an important role in cell division of yeast cells in macrophages and affect asexual development. *In vitro* overexpression of *pakB* alleles with either a mutated CRIB (<u>C</u>dc42/<u>R</u>ac Interactive <u>B</u>inding) or GBB (<u>Gβ</u> <u>b</u>inding) domains, results in the production of yeast cells that resembled those produced during growth inside macrophages (Boyce et al. 2009). Additionally MsgA and PakB have complementary localisation patterns at septation sites during intracellular yeast formation. Together these are suggestive of coordinated effects on morphogenetic mechanisms that involve the regulation of cell division. Similar localization and morphological function has been observed in *S*. *cerevisiae* for the GEF-like protein Lte1, which plays a vital role in activating the Mitotic Exit Network (MEN) and the formation of daughter bud cells. Lte1 is localized to the incipient bud cortex and *lte1* mutants show delayed cytokinesis and aberrant daughter cell morphology (Bertazzi et al. 2011; Geymonat et al. 2009). Lte1 is activated by the PakB homologue Cla4, which enables its localisation to the bud where it inhibits machinery that prevents mitotic exit and thus proper bud formation (Höfken & Schiebel 2002; Jensen et al. 2002; Seshan et al. 2002; Yoshida et al. 2003). In this respect LTE1 does not function as a GEF but rather in a separate signalling pathway controlling morphogenesis. *T*. *marneffei* and other filamentous fungi lack a clear, conserved orthologues of Lte1 and components involved in MEN, suggesting that they use an alternate pathway to accomplish this role and this may include the MsgA and PakB.

### Unique domain structure of MsgA influences its function and localisation

The predicted gene product of *msgA* possesses a DH domain stereotypical of GEFs and involved in regulating GTP dependent interactions. However it lacks the auxiliary PH domain that is common to canonical GEFs and important for localisation. Instead MsgA has a BAR domain that is not present in any of the other Ras, Rho, Rsr, Arf, Rab and Ran GEFs in *T*. *marneffei*, but which is conserved in orthologues of MsgA in other ascomycete fungi. The function of the BAR domain has been characterised in a number of systems and it is clear that it is involved in membrane dynamics, particularly by its ability to oligomerise and induce membrane bending, but can also bind small GTPases and affect their function (Habermann 2004). The data shows that a *msgA* allele missing the BAR domain (*msgA*^ΔBAR^) failed to complement the aberrant yeast morphology phenotype observed in the Δ*msgA* strain whereas a *msgA* allele missing the DH domain (*msgA*^ΔDH^) fully rescued this phenotype. At least part of basis for this lack of complementation is likely to be due to the resultant loss of localisation of MsgA when the BAR domain is removed. This hypothesis is consistent with the correct localisation, and complementation, of the MsgA mutant protein that contains the BAR domain but lacks the DH domain. Like a number of other BAR domain proteins this suggests that part of its role may reside in its interaction with, and recruitment of, other factors to specific sites on the cell membrane (Miki et al. 2000; Schorey et al. 1997).

Proteins containing BAR domains are known to interact with many small GTPases including Rac (Tarricone et al. 2001; Van Aelst et al. 1996; Schorey et al. 1997). In *T*. *marneffei* the Rac- (*cflB*), Cdc42- (*cflA*) and Ras- encoding (*rasA*) genes have been shown to play important and distinct roles in hyphal and yeast morphogenesis *in vitro*, yeast morphogenesis *ex vivo*, conidial germination and asexual development (Boyce et al. 2001; Boyce et al. 2003; Boyce et al. 2005), yet none of the phenotypes associated with the various mutant alleles examined in these studies, which include deletion, dominant activating and negative alleles, mimic those for *msgA* alleles. The *rasA*, *cflA* and *msgA* genes play a role in yeast cell morphogenesis but the *msgA* effects are restricted to *ex vivo* yeast while the *rasA* and *cflA* effects occur in all yeast cells. Moreover, the phenotypes of the *msgA* and *rasA*/*cflA* mutants with respect to yeast morphology are strikingly different. Similarly, the *cflB* and *msgA* genes play a role in asexual development but the effects of perturbing their function are different. Coupled with the observation that deletion of the DH domain does not appear to affect MsgA function and that *msgA* mutant yeast cells do not express *in vitro* yeast cell- specific genes, it seems that *msgA* controls yeast cell morphogenesis independently of these small GTPases and does not function in host recognition and signalling.

Canonical Rho GEFs have been investigated in a number of pathogenic fungi, and shown to play roles in cellular morphogenesis and pathogenicity (Wendland & Philippsen 2001; Bassilana et al. 2003; Fuchs et al. 2007; Tang et al. 2005). Similarly, non-GEF-like BAR domain proteins have been associated with morphology and virulence in a number of plant and human pathogenic fungi. For example, *C*. *albicans rvs* mutants affect hyphal morphogenesis, invasive growth and antifungal resistance through their involvement in endocytic mechanisms (Douglas et al. 2009). Also the rice blast fungus *M*. oryzae BAR domain proteins Rvs161 and Rvs167 are important for plant invasion through appresorial formation (Dagdas et al. 2012). However non-canonical GEF-like proteins with a BAR domain are poorly understood in most systems, including fungi. The best example of a DH domain protein with an associated BAR domain is the human dynamin-binding scaffold protein Tuba, which localizes to brain synapses as puncta at the base of membrane ruffles and is involved in synaptic vessel endocytosis (Salazar et al. 2003; Kovacs et al. 2006). Membrane ruffling plays a crucial role in internalization during substrate acquisition and receptor availability control as well as cell motility (Hoon et al. 2012). Deletion of the BAR domain in this Tuba abolishes dorsal ruffling of synaptic membranes and causes the mislocalisation of this protein throughout the cytoplasm, suggesting that the BAR domain is necessary for facilitating correct localisation of Tuba during synaptic cell cytoskeletal dynamics (Kovacs et al. 2006). The phenotypic similarities between MsgA and this distantly related mammalian protein are striking and support the idea that BAR domains are important factors in determining cellular morphology across kingdoms.

Analysis of the *msgA* gene of *T*. *marneffei* shows that it is important for yeast cell morphogenesis during macrophage infection and in conidial production during asexual development. Although yeast cells divide by fission and conidia are formed by budding, both of these cell types require cell separation, which is unlike hyphal cells. Yeast cells formed *in vitro* also require cell separation but *msgA* does not appear to be required for this process. The distinction here may lie in the fact that yeast cell morphogenesis *in vitro* is always preceded by hyphal growth and the process of arthroconidiation that produces yeast cells *in vitro* may be mechanistically distinct. The data presented here suggests that *msgA* and the p21-activated kinase encoding *pakB* may function in the same pathway to control morphogenesis during infection. Further studies to identify binding partners of MsgA will shed light on the mechanism that controls cellular morphogenesis of *T*. *marneffei* during infectious growth.

## Supporting information

