## Supplementary material for "The novel *Dbl* homology/BAR domain protein, MsgA, of *Talaromyces marneffei* regulates yeast morphogenesis during growth inside host cells"

**Supplementary Data**

**Supplementary Figure 1: Unique protein structural domain composition and sequence features of the MsgA protein.**

The MsgA protein has unique structural domain that differs from other canonical RhoGEF proteins. Additionally sequence alignment of the MsgA protein reveals a region containing tandem glutamic acid repeats, showing variation in number amongst dimorphic pathogens, *Talaromyces* clade and clinical isolates (A) Schematic of MsgA protein highlighting the specific domain structure, which includes the Dbl homology domain (DH, blue) and Bin-Amphiphysin-Rvs domain (BAR, green). Vertical black bar within the N-terminal domain represents the 26 glutamic acid repeat string. Numbers represent amino acid residues in MsgA protein. Schematics of protein domain organisation for the remaining 5 RhoGEF proteins highlight the unique structure of the MsgA protein. Other domains represented include plekstrin homology domain (PH, purple), Cdc42/Scd1 domain (Cdc42/Scd1, grey) Phox and Bem1 interacting domain (PB1, yellow), Dishevelled, Egl-10 and Pleckstrin domain (DEP, brown) and Citron homology domain (CNH, green). (B) Sequence alignment of the tandemly repeated glutamic acid region of the MsgA protein for *Talaromyces funiculosus* (Tf), *Talaromyces flavus* (Tfl), *Talaromyces stipitatus* (Ts) (above dashed line) and 14 isolates of *T. marneffei* (below dashed line). While the non-pathogenic closest fungal relatives *Tf*, *Tfl* and *Ts* show the smallest number of glutamic acid repeats, each of the clinical isolates show a varying number of repeats ranging from 15 - 26. The *T. marneffei* isolates include the standard clinical isolates from Hong Kong (Pm1) and Thailand (F4), a collection from AIDS patients (3482, 3840, 3041, 3871, 4059, 027 and 043), a number of patients without AIDS (012, 203 and 702) and isolates from the Bamboo rat (*Rhizomys sinensis*, a natural host of *T. marneffei*)(HR2 and BR2). Sequences for *T. funiculosus*, *T. flavus* and 13 clinical and non-clinical isolate strains (3482, 3840, 3041, 3871, 4059, 027, 043, 012, 203, 702, HR2, BR2 and F4) were derived from a genome sequencing study (Payne *et al* in prep). (C) Sequence alignment of the glutamic acid region (grey) for *T. marneffei* and other non-pathogenic and pathogenic fungal species. Tandem repeats of glutamic acid are seen only in the *T. marneffei* orthologue of MsgA. The *T. marneffei* (Tm 2161) sequence used in the alignments is the type strain. These include non-pathogenic *Aspergillus nidulans* (An), pathogenic *Aspergillus fumigatus* (Af) and dimorphic pathogens *Coccidioides immitis* (Ci), *Histoplasma capsulatum* (Hc) and *Paracoccidioides* *brasiliensis* (Pb). The (*) indicates positions which have a single, fully conserved residue, (:) indicates conservation between residues that have strongly similarities and (.) indicates conservation between residues that have weak similarities. Alignments were done using CLUSTAL O (1.2.0).

**Supplementary Figure 2: The filaments produced by the *ΔmsgA* strain do not exhibit transcriptional patterns characteristic of 37˚C *in vitro* arthroconidia or yeast cells.**

Expression of genes with 37˚C *in vitro* specific expression (represented by PMAA accession IDs) in the wildtype and Δ*msgA* strains, during both *in vitro* and intracellular macrophage growth were used to determine if the aberrant filaments observed in the *ΔmsgA* mutant strain were similar to 37˚C *in vitro* growth. RNA isolated from wildtype grown for 6 days in liquid BHI medium (WT *in vitro*), ΔmsgA strain grown in LPS activated J774 murine macrophages for 24 hours *(ΔmsgA* intracellular) and wildtype grown in LPS activated J774 murine macrophages for 24 hours (WT intracellular). The histone H3 gene was used as a control (H3). The *ΔmsgA* strain does not show inappropriate expression of these genes during macrophage infection and is comparable to expression patterns seen in wildtype isolated from within macrophages, suggesting that the filaments and elongated yeast cells observed during macrophage infection of this strain are not similar to 37˚C *in vitro* grown yeast.

**Supplementary Figure 3: The *ΔmsgA* strain produces elongated yeast cells in THP-1 macrophages at 24 hours post infection.**

Differentiated THP-1 human macrophages were infected with conidia from the wildtype, Δ*msgA* and complemented Δ*msgA msgA*$^+$ strains and examined microscopically after 24 hours. (A) At 24 h post infection wildtype (*msgA*$^+$) *T. marneffei* within THP-1 macrophages produced numerous ovoid yeast cells that divided by fission. In contrast the *ΔmsgA* mutants show increased yeast cell length. The complemented strain (Δ*msgA msgA*$^+$) produced yeast cells of comparable to wildtype. (B) The effect of deleting *msgA* on morphogenesis in macrophages was quantified at the 24 h post infection time point. Wildtype yeast cells showed an average length of 4.9±0.09μm compared to the Δ*msgA* mutant strain, which produced yeast cells of an average cells length of 8.6±0.25μm, approximately 1.7 times longer than wildtype. Images were captured using differential interference contrast (DIC) or with epifluorescence to observe fungal cell walls stained with calcofluor white (CAL). Error bars represent the standard error of the mean with t-test values falling in the following range *** ≤0.001. Scale bars are 10 μm.

**Supplementary Figure 4: Continuous overexpression of *msgA* at 37°C *in vitro* causes loss of polarisation and increases pigment production**

The wildtype (*msgA+*) and *xylP(p)::msgA* strains were grown for 6 days in BHI medium either with 1% xylose (induced) or without supplementation (uninduced). This growth without transfer leads to continuous induction. (A) Under continuous induction wildtype *T. marneffei* produces elongated filament like yeast cells. However the *xylP(p)::msgA* strain produces swollen arthroconidia (single arrowhead) and chains of yeast cells that resembles growth during intracellular macrophage infection. Additionally continuous induction of the *xylP(p)::msgA* strain produces yeast cells that have polarity defects which result in the formation of multiple yeast cells from the poles (single arrowhead), unlike wildtype where daughter cells are produced via fission driven separation from either pole of the primary yeast cell (double arrowhead). Images were captured using differential interference contrast (DIC) or with epifluorescence to observe fungal cell walls stained with calcofluor white (CAL). Error bars represent the standard error of the mean with t-test values falling in the following range *** ≤0.001. Scale bars are 10 μm.

**Supplementary Figure 5: The *ΔmsgA* mutant strain is involved in conidial production during asexual development**

Wildtype (*msgA+*). *ΔmsgA*, *ΔmsgAmsgA$^+$* and *xylP(p)::msgA* strains were grown for 10 days in ANM medium with 1% glucose (Uninduced) or 1% xylose (Induced). (A) Colonial appearance of each strain grown in non-inducing and inducing media reveal differences in pigmentation. Notably at the *ΔmsgA* mutant strain appears paler at the colonial level compared to wildtype. (B) In both conditions the *ΔmsgA* mutant strain showed reduced conidial production compared to wildtype. The *ΔmsgA* mutant strain showed reduced conidiation both with and without induction compared to wildtype ($5.3 \times 10^8 \pm 0.4$ vs. $23.9 \times 10^8 \pm 0.9$ conidia/ml and $6.3 \times 10^8 \pm 0.9$ vs. $33.9 \times 10^8 \pm 1.5$ conidia/ml respectively). This phenotype was not completely rescued by the reintroduction of the wildtype allele in the *ΔmsgA msgA$^+$* strain ($15.9 \times 10^8 \pm 1.1$ conidia/ml without induction and

18.5x10$^{8}$±1.3 conidia/ml with induction). The *xylP(p)::msgA* strain mimicked the *ΔmsgA* mutant strain during growth without induction (7.7x10$^{8}$±0.8 conidia/ml). However under inducing conditions the *xylP(p)::msgA* strain showed complete restoration of conidiation to wildtype levels (35.2x10$^{8}$±1.5 conidia/ml). Error bars represented standard error of the mean with t-test values falling in the following range *** ≤0.001.

**Supplementary Table 1: Primers used in this study.**

| Primer ID | Primer sequences | Origin |
|---|---|---|
| VV55 | attgccatagagaccgct | This study |
| VV56 | tgtagaactcccccaggt | This study |
| VV57 | ggggacccagctttcttgtacaaagtggtaaggagacagcatagcac | This study |
| VV58 | ggggagcctgctttttttgtacaaacttgtgaaggactgaagcggtta | This study |
| WW42 | aggatgtaaccgttgagg | This study |
| WW44 | caacaaaaactgggacctatt | This study |
| WW45 | tgtgacctacgatgattagca | This study |
| WW46 | tttttctagaaccaagctttgcaaaatgctgt | This study |
| WW57 | caattcactggccgtcgtttt | This study |
| WW59 | gtggtcggtcaagttgttagc | This study |
| WW60 | ctgttctggtgttggtgagcc | This study |
| WW77 | ccgttggcgaggatggtgata | This study |
| WW78 | agaaacaaacctggctaccct | This study |
| WW79 | aaaccaccgaagtctatccaa | This study |
| H57 | tgtaaaacgacggccagt | (Boyce et al., 2010) |
| H56 | ggaaacagctatgaccatg | (Boyce et al., 2010) |

**Supplementary Table 2. *T. marneffei* strains used in this study.**

| Strain ID | Full Genotype | Origin |
|---|---|---|
| G809 | *ΔligD::pyrG*$^{+}$*niaD pyrG* | Bugeja *et al.*, 2012 |
| G816 | *ΔligD niaD*1 *pyrG* | Bugeja *et al.*, 2012 |
| G829 | *ΔligD ΔriboB::pyrG*$^{+}$ *niaD pyrG* | Bugeja *et al.*, 2012 |
| G1044 | *ΔligD niaD pyrG ΔmsgA::pyrG*$^{+}$ | This study |
| G1045 | *ΔligD niaD pyrG ΔmsgA::pyrG*$^{+}$ [niaD$^{t}$ *msgA*$^{+}$] | This study |
| G1046 | *ΔligD niaD pyrG ΔmsgA::pyrG*$^{+}$ [niaD$^{t}$ *msgA*$^{+}$::mCherry] | This study |
| G1047 | *ΔligD niaD pyrG ΔmsgA::pyrG*$^{+}$ [niaD$^{t}$ *msgA*$^{ΔBAR}$] | This study |
| G1048 | *ΔligD niaD pyrG ΔmsgA::pyrG*$^{+}$ [niaD$^{t}$ *msgA*$^{ΔDH}$] | This study |
| G1049 | *ΔligD niaD pyrG ΔmsgA::pyrG*$^{+}$ [niaD$^{t}$ *msgA*$^{ΔBAR}$*::mCherry*] | This study |
| G1050 | *ΔligD niaD pyrG ΔmsgA::pyrG*$^{+}$ [niaD$^{t}$ *msgA*$^{ΔDH}$*::mCherry*] | This study |
| G1051 | *ΔligD niaD pyrG*$^{+}$ *xylP(p)::msgA::bar*$^{+}$ | This study |

**Supplementary Table 3. Kinetics of conidial germination during *in vitro* growth and macrophage infection.**

| ***In vitro*** | | | |
|---|---|---|---|
| **Strain** | **16 hours** | **24 hours** | **48 hours** |
| *msgA*$^+$ | 59±2.47 | 89.71±3.06 | 98.1±2.62 |
| Δ*msgA* | 53.8±0.49 | 86.71±1.21 | 96.04±0.76 |
| **Intracellular growth** | | | |
| **Strain** | **4 hours** | **6 hours** | |
| *msgA*$^+$ | 38.8±3.5 | 72.7±9.5 | |
| Δ*msgA* | 37.1±2 | 75.3±5 | |

**A**

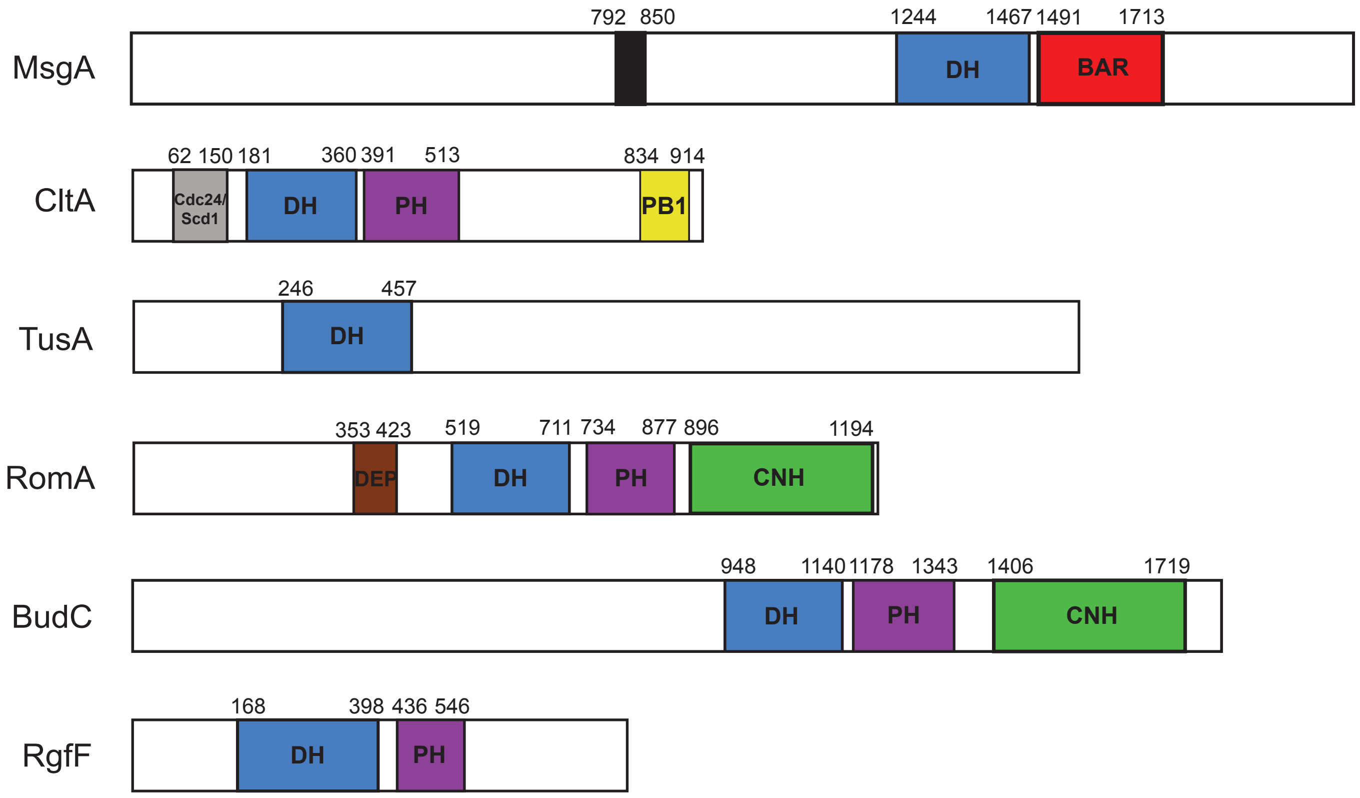


**B**

```
Tf      769   ASHRDLLSRIMQLRESSSSSECDEEEED----------------------------DDDD
Tfl     788   ASHRDLLSRIMQLRESSSSSECDEEEED----------------------------DDDN
Ts      788   ESHRDLLSRIMQLRESSSSSECDEDED-----------------------------DDDD
.............................................................................
2161    780   ASHRDLLSRIMQLRESSSSSECDEEEEEEEEEEEEEEEEEEEEEEEEEEDDDDDDEDEDD
Pm1     780   ASHRDLLSRIMQLRESSSSSECDEEEEEEEEE---------EEEEEEEEEEDDDDDDEDD
F4      780   ASHRDLLSRIMQLRESSSSSECDEEEEEEEEE----------EEEEEEEDDDDDDEDEDD
3482    780   ASHRDLLSRIMQLRESSSSSECDEEEEEEEEE-----------EEEEEEEEDDDDDDEDD
3840    727   ASHRDLLSRIMQLRESSSSSECDEEEEEEEEE---------EEEEEEEEDDDDDDEDEDD
3841    780   ASHRDLLSRIMQLRESSSSSECDEEEEEEEEEEEE---EEEEEEEEEEDDDDDDDEDEDD
3871    780   ASHRDLLSRIMQLRESSSSSECDEEEEEEEEE-----------EEEEEEEEDDDDDDEDD
4059    780   ASHRDLLSRIMQLRESSSSSECDEEEEEEEEE---------EEEEEEEEDDDDDDEDEDD
027     780   ASHRDLLSRIMQLRESSSSSECDEEEEEEEEE-----------EEEEEDDDDDDDEDEDD
043     780   ASHRDLLSRIMQLRESSSSSECDEEEEEEEEE----------EEEEEEDDDDDDDEDEDD
012     780   ASHRDLLSRIMQLRESSSSSECDEEEEEEEEE-------------EEEEDDDDDDEDEDD
203     780   ASHRDLLSRIMQLRESSSSSECDEEEEEEEEE-----------EEEEEDDDDDDDEDEDD
702     780   ASHRDLLSRIMQLRESSSSSECDEEEEEEEEE-----------EEEEEEDDDDDDEDEDD
HR2     780   ASHRDLLSRIMQLRESSSSSECDEEEEEEEEE-------------EEEEEEDDDDDDEDD
BR2     780   ASHRDLLSRIMQLRESSSSSECDEEEEEEEEE-----------EEEEEDDDDDDDEDEDD
               ***********************:*:                             *:*:
```

**C**

```
Tm 2161   780   ASHRDLLSRIM--------QLRESSSSSE--CDEEEEEEEEEEEEEEEEEE---EEEEEEEEDDDDDDEDEDDNELCDDEND
An        753   TSHRTLLNEIM--------RIRESSPSSSS-CDEQE---------------------------------------YGFSDND
Af        758   ASHRTLLSQIM--------QIRESSPDTSS-CDEAD---------------------------------------YSFSDTE
Ci        722   QSHRSMLSHIM--------QMRESSPDITDSSDDDE---------------------------------------DTSSSGR
Hc        760   QIHRTVLSQIM--------QMRESSPSSSSSSD-------------------------------------------SFSERD
Pb1       761   QIHRTVLSQIM--------QMRESSPSSSAYSD-------------------------------------------SFSERD
Pb2       268   TSIRILRSRLAQPV----RTIVQSEPTSVDSHEFRA---RIRK---------------------------------KLNPSI
                     :              : : .       :
```

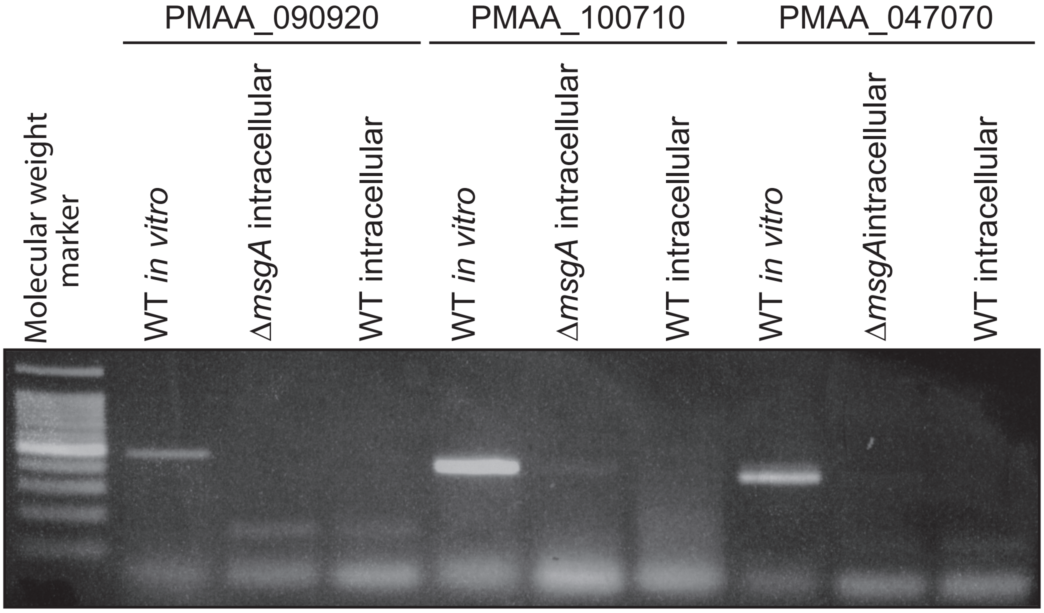
PMAA_090920
PMAA_100710
PMAA_047070
Molecular weight marker
WT *in vitro*
ΔmsgA intracellular
WT intracellular
WT *in vitro*
ΔmsgA intracellular
WT intracellular
WT *in vitro*
ΔmsgAintracellular
WT intracellular

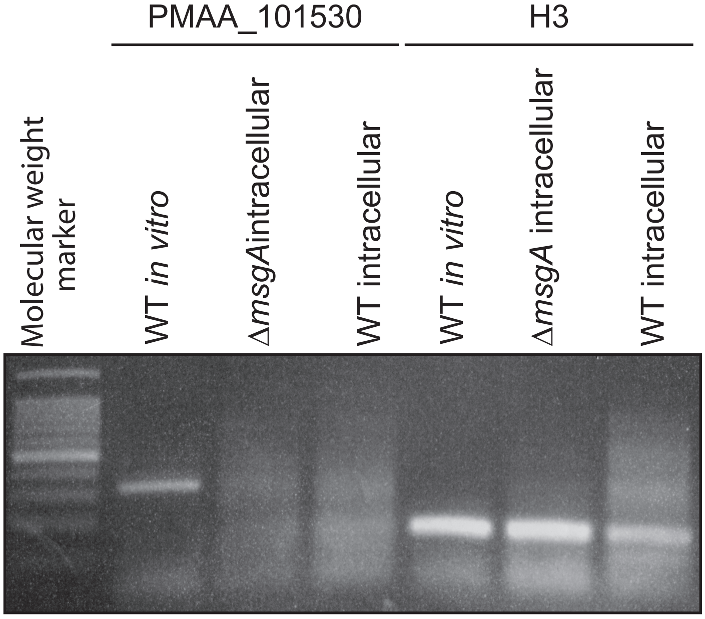
PMAA_101530
H3
Molecular weight marker
WT *in vitro*
ΔmsgAintracellular
WT intracellular
WT *in vitro*
ΔmsgA intracellular
WT intracellular

**A**

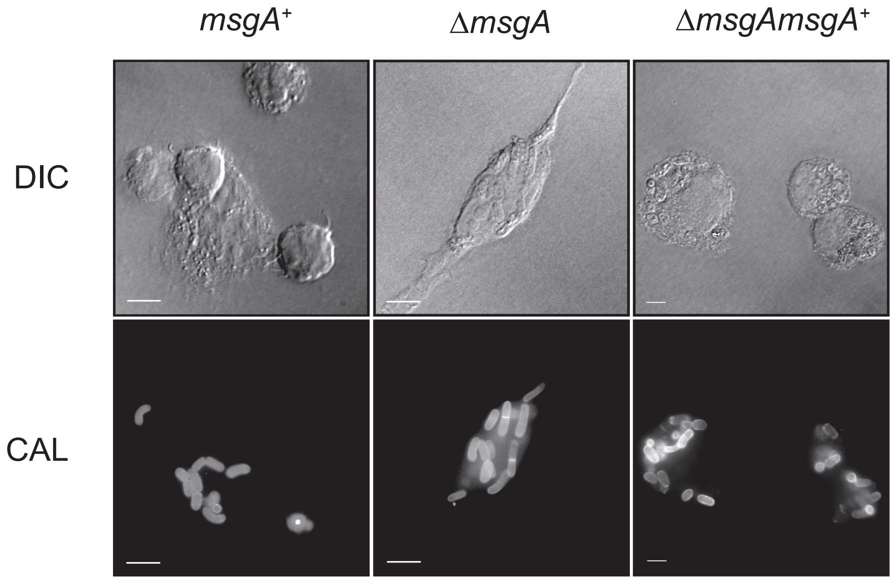
msgA+
ΔmsgA
ΔmsgAmsgA+
DIC
CAL

**B**

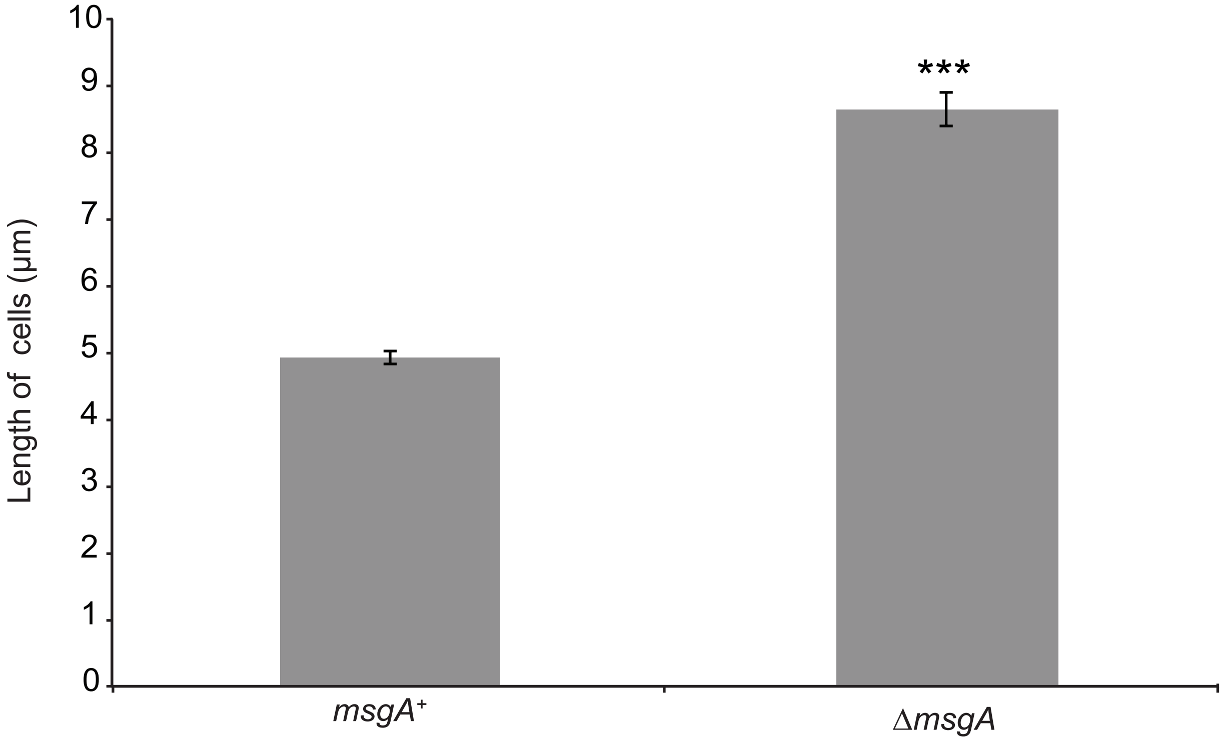
***
Length of cells (µm)
0
1
2
3
4
5
6
7
8
9
10
msgA+
ΔmsgA

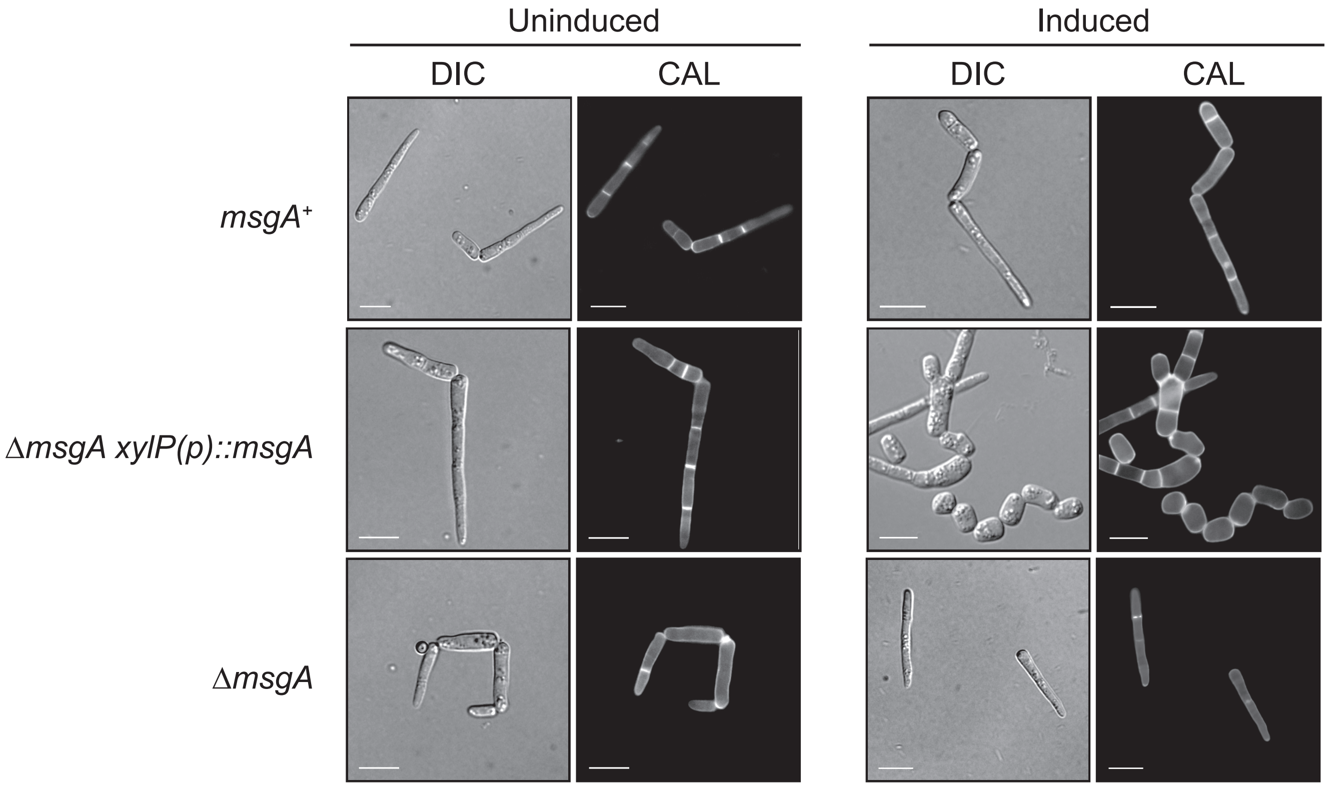

Uninduced
Induced
DIC
CAL
DIC
CAL
msgA+
ΔmsgA xylP(p)::msgA
ΔmsgA

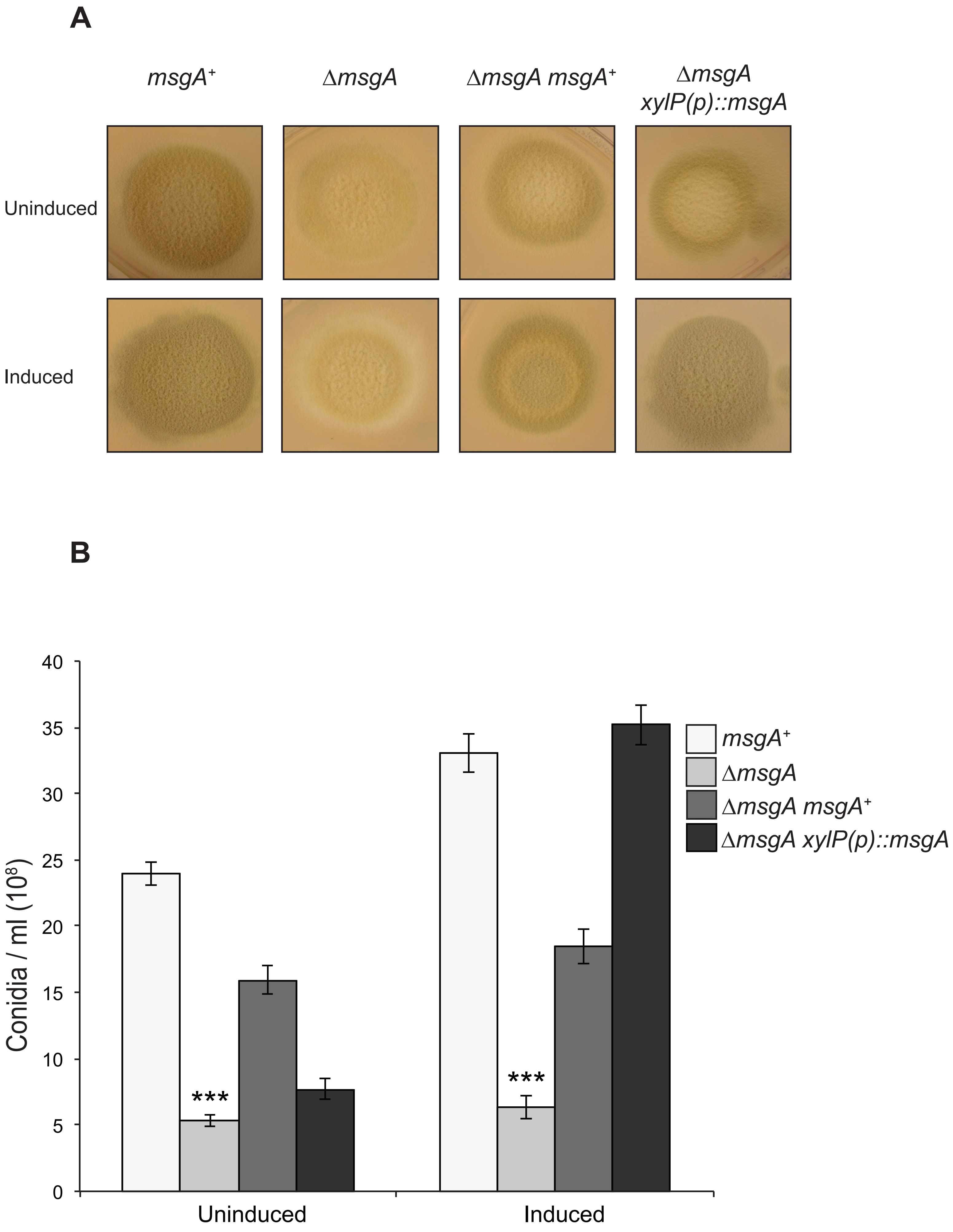
A
msgA+
ΔmsgA
ΔmsgA msgA+
ΔmsgA xylP(p)::msgA
Uninduced
Induced
B
Conidia / ml (10^8)
40
35
30
25
20
15
10
5
0
***
***
Uninduced
Induced
msgA+
ΔmsgA
ΔmsgA msgA+
ΔmsgA xylP(p)::msgA
